# The city and forest bird flock together in a common garden: genetic and environmental effects drive urban phenotypic divergence

**DOI:** 10.1101/2024.08.27.609854

**Authors:** M.J. Thompson, D. Réale, B. Chenet, S. Delaitre, A. Fargevieille, M. Romans, S.P. Caro, A. Charmantier

## Abstract

Urban phenotypic divergences are documented across diverse taxa, but the underlying genetic and environmental drivers behind these phenotypic changes are unknown in most wild urban systems. We conduct a common garden experiment using great tit (*Parus major*) eggs collected along an urbanization gradient to: 1) determine whether documented morphological, physiological, and behavioural shifts in wild urban great tits are maintained in birds from urban and forest origins reared in a common garden (N = 73) and 2) evaluate how different sources of genetic, early maternal investment, and later environmental variation contributed to trait variation in the experiment. In line with the phenotypic divergence in the wild, common garden birds from urban origins had faster breath rates (i.e., higher stress response) and were smaller than birds from forest origins, while wild differences in aggression and exploration were not maintained in the experiment. Differences between individuals (genetic and environmentally induced) explained the most trait variation, while variation among foster nests and captive social groups was limited. Our results provide trait-specific evidence of evolution in an urban species where genetic change likely underlies urban differences in morphology and stress physiology, but that urban behavioural divergences are more strongly driven by plasticity.

## Introduction

Various evolutionary and ecological processes shape the phenotypic diversity we observe in nature and lead to phenotypic divergences between populations (Mitchell-Olds et al. 2007; Hendry 2017). For instance, local selection pressures on heritable traits can lead to divergent adaptation to local environmental conditions (Kawecki and Ebert 2004). Developmental or reversible phenotypically plastic responses to local environments can also drive phenotypic adjustments (Ghalambor et al. 2007). Determining how local adaptation and plasticity interact to shape phenotypes is crucial as these processes can have different impacts on demographic and evolutionary trajectories of wild populations (Ghalambor et al. 2007; Snell-Rood 2013; Nicolaus and Edelaar 2018).

Some of the most striking examples of phenotypic diversity occur along urbanization gradients, such as urban shifts in multiple taxa towards smaller body sizes (Merckx et al. 2018). Such phenotypic shifts in urban populations are frequently documented across diverse taxa and traits (Szulkin et al. 2020; Diamond and Martin 2021), through changes in both phenotypic means (Miranda et al. 2013; Lambert et al. 2021) and more recently in phenotypic variation (Capilla-Lasheras et al. 2022; Thompson et al. 2022). Urban phenotypic divergences are commonly assumed to be driven by genetic change via selection, but there is still a lack of evidence that urban organisms are adapting to these novel urban conditions (Lambert et al. 2021) and plasticity could play a major role in urban phenotypic change (Yeh and Price 2004; Hendry et al. 2008). Determining the mechanisms behind phenotypic changes in urban organisms could importantly inform on whether urban populations will continue to adjust in pace with further environmental change. For these reasons, there have been several calls for research that disentangle the genetic and plastic contributions on urban phenotypes and, more specifically, calls for urban common garden experiments (Rivkin et al. 2019; Lambert et al. 2021; Sanderson et al. 2023).

Common garden experiments are a useful approach for exploring the genetic basis of phenotypic differences between populations. These manipulations rear individuals from different populations under the same environmental conditions from very early life stages, and ideally across several generations. As individuals develop and mature under common conditions, phenotypic differences that persist in this context should reflect underlying genetic differentiations rather than plastic responses to environmental conditions (Lambert et al. 2021). Thus, common gardens can help determine whether evolutionary (i.e., genetic) change drives documented phenotypic divergences in wild populations, and can be used to explore interactions between genetic and plastic changes acting in these systems (Conover et al. 2009). For these reasons, common garden approaches are needed in urban evolution research to explore the potential processes acting in these contexts (Alberti et al. 2017; Sanderson et al. 2023).

We censused 77 common garden studies with urban populations in the literature; an impressive number despite the effort and resources these experiments require (see synthesis in Table 1). Most of these studies have been published within the last ten years (83%, N = 64) and many support genetic divergence underlying shifts in urban phenotypes (86%, N = 66), which could indicate local adaptation to urban conditions via evolution. Fewer studies document plasticity to environmental conditions as a driver of phenotypic change (58%, N = 45), but this conclusion is especially common in multi-treatment common gardens where individuals are reared under multiple environmental treatments (e.g., temperature treatments). Experiments so far tend to use invertebrate or plant models (75%, N = 58), likely as these organisms are more easily reared, reproduced, and manipulated in captive environments. Many studies have focused on physiological phenotypes associated with tolerance to temperature as the urban heat island effect is known to increase temperature and heat stress in urban environments (Mohajerani et al. 2017). For example, urban damselflies (*Coenagrion puella*), water fleas (*Daphnia magna*), wood louse (*Oniscus asellus*), and acorn ants (*Temnothorax curvispinosus*) have higher heat tolerance compared to nonurban conspecifics when reared under common conditions, providing evidence that urban invertebrates have adapted to urban heat islands (Brans et al. 2017; Diamond et al. 2018; Tüzün and Stoks 2021; Yilmaz et al. 2021).

**Table 1.**
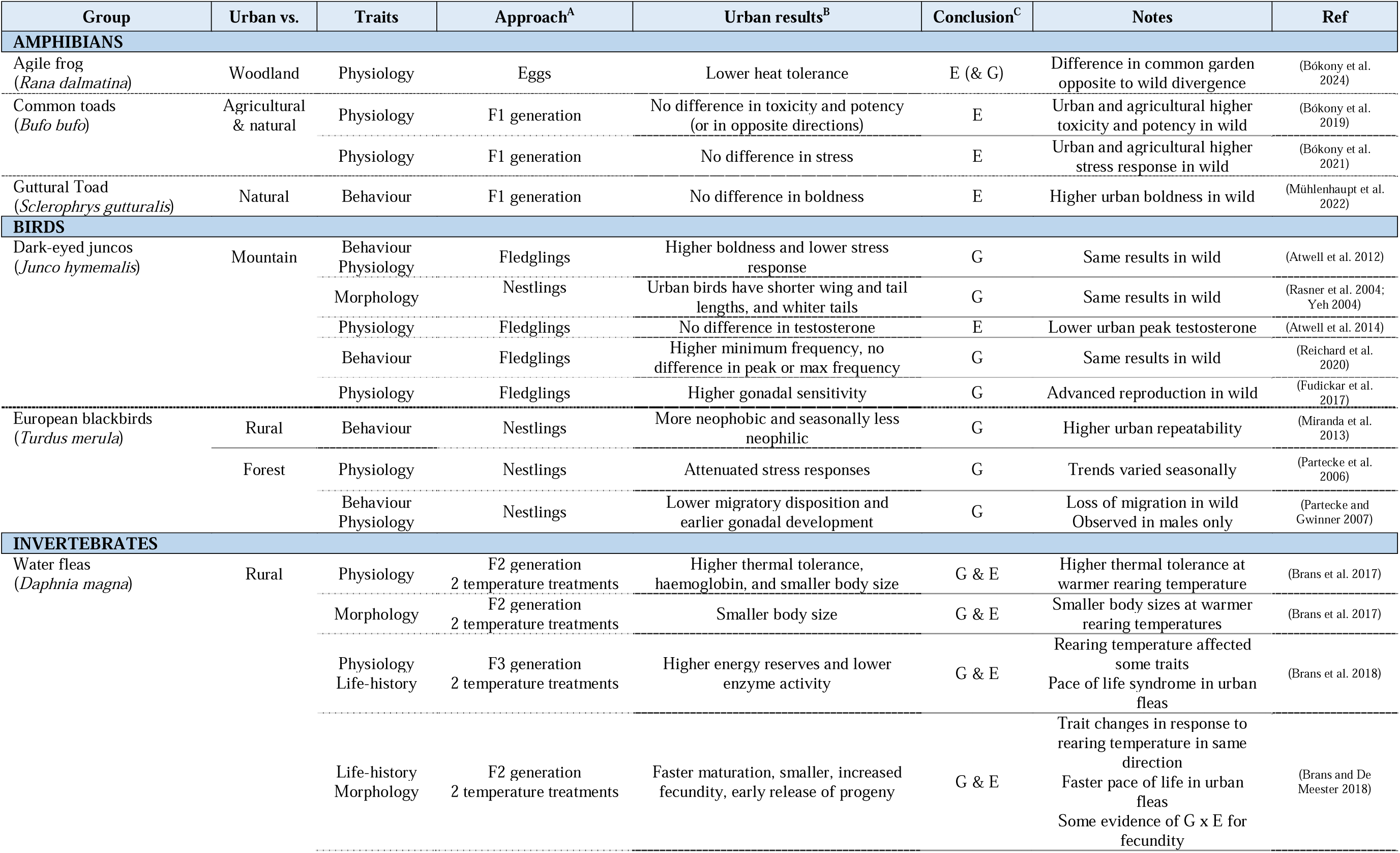
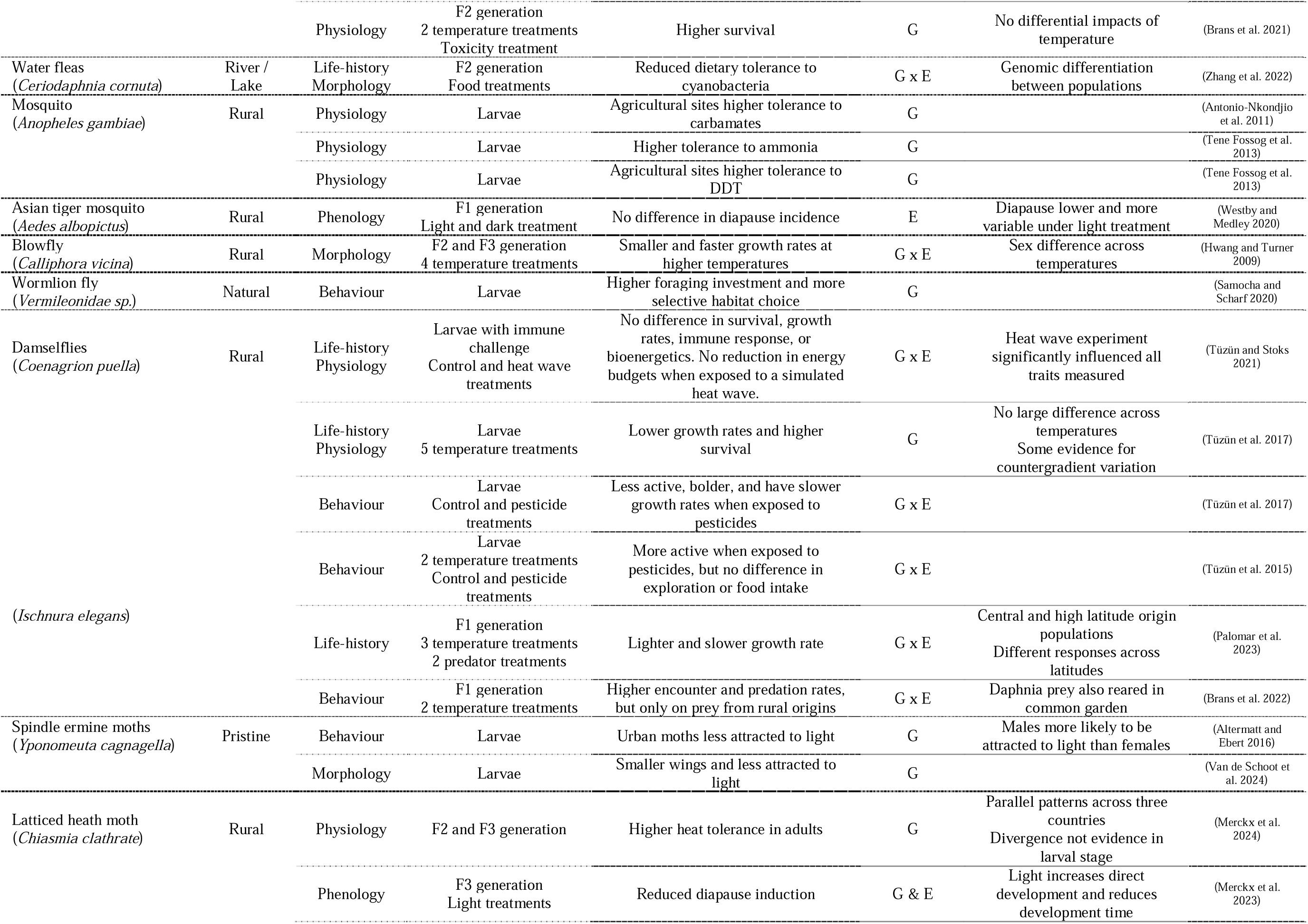
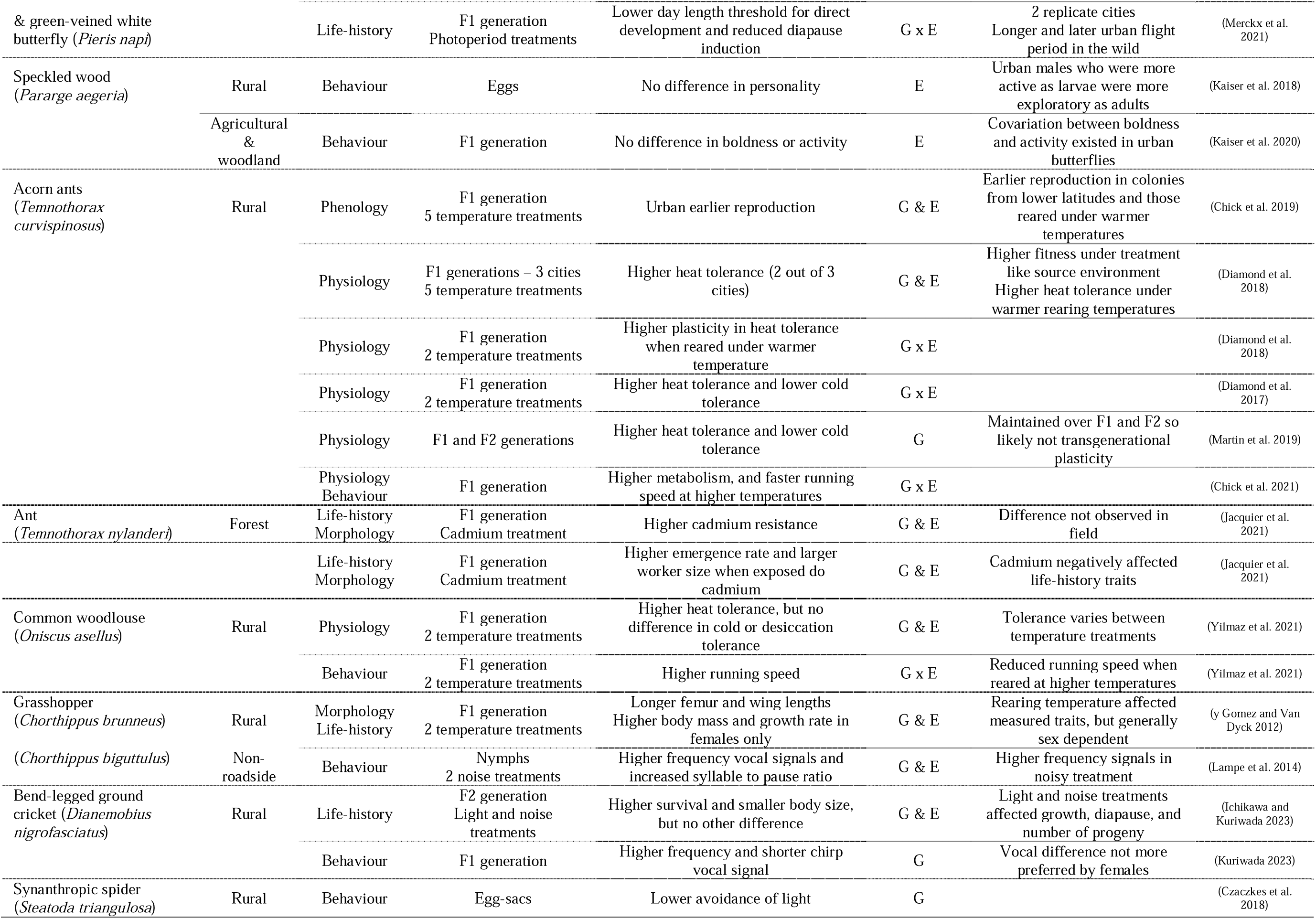
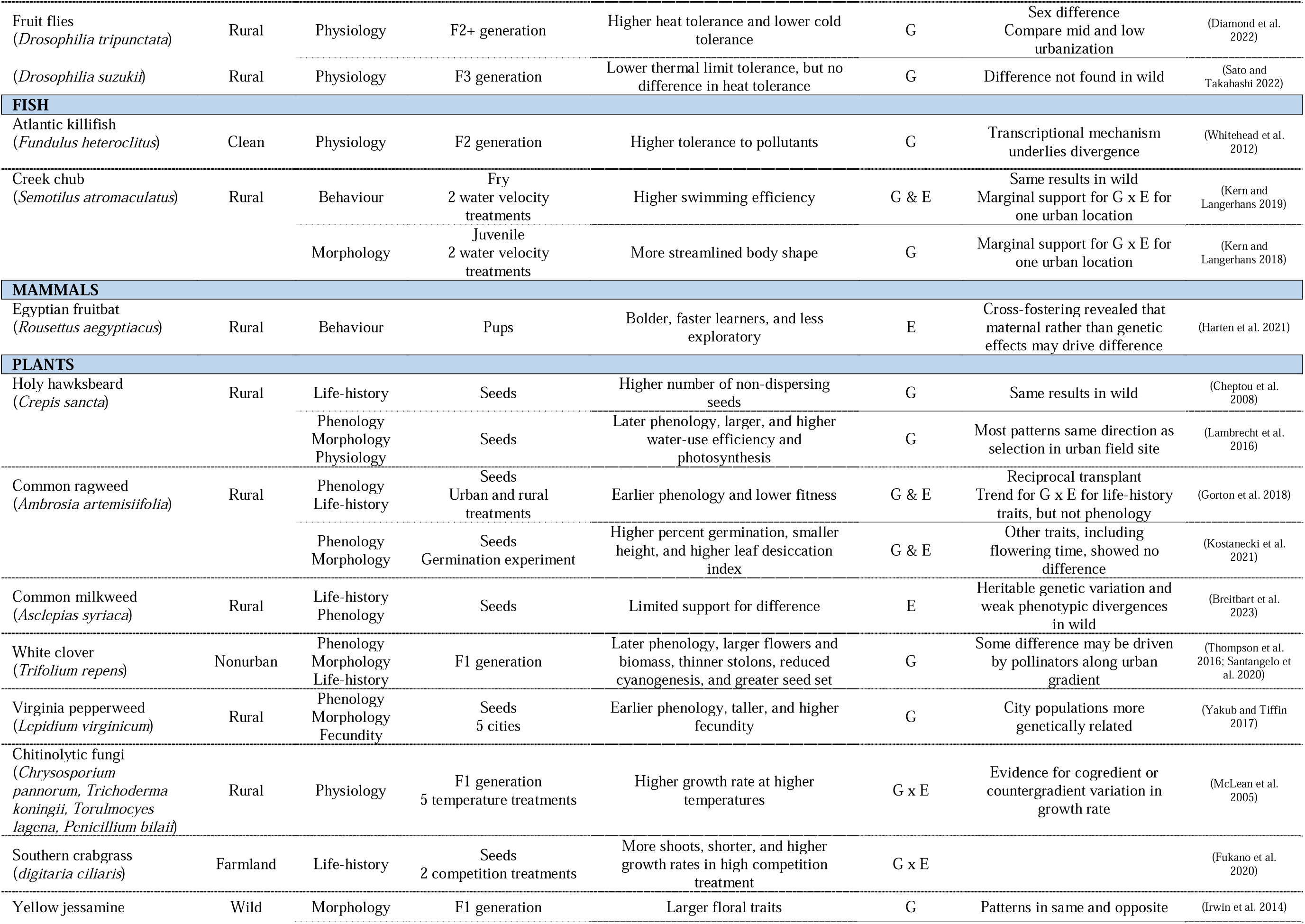
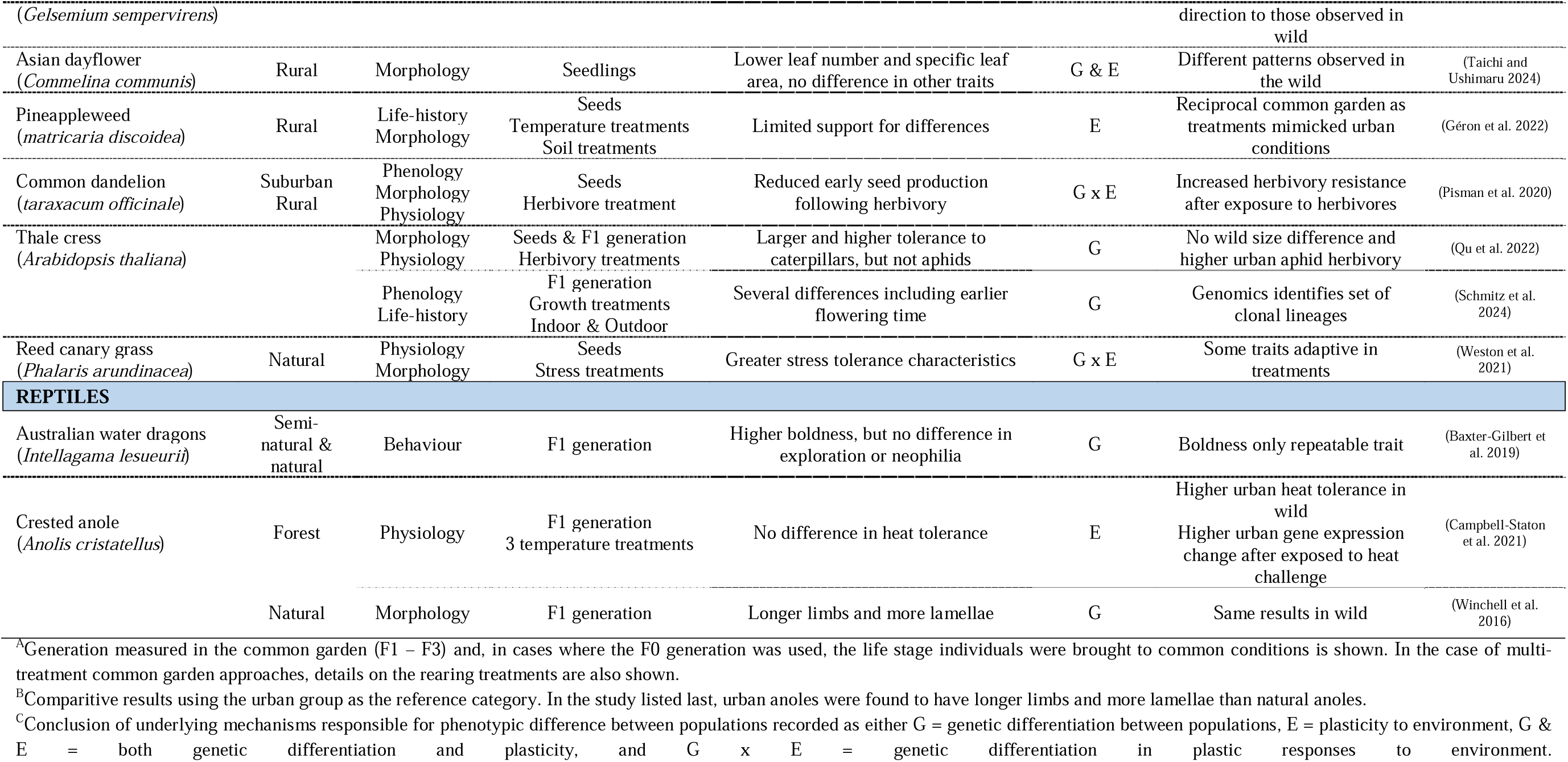
Synthesis of studies (*N* = 77) that have used a common garden approach to compare the phenotypes of urban and nonurban populations (e.g., rural, forest, mountain, agricultural) across a variety of different groups (amphibians, birds, invertebrates, fish, plants, reptiles) and traits (physiology, behaviour, morphology, life history, phenology). Information concerning the traits measured, approach taken, results, conclusions, and other notes are shown for each study. Studies were collated from Lambert et al. 2021 (Table 1) and from a literature search using Google Scholar for articles since 2020 that included both “common garden” and “urban” (conducted March 22 2024).

While common garden studies provide support in favour of urban evolution in some well-studied invertebrate and plant taxa, whether genetic change drives phenotypic shifts in other urban taxa is not as well known (e.g., Table 1: 10% or N = 8 common garden studies in birds). For instance, birds are one of the most studied taxa in urban ecology and evolution and there are growing generalizations on how urbanization impacts the traits of birds globally (e.g., earlier lay dates or smaller body sizes, Capilla-Lasheras et al. 2022; Hahs et al. 2023). Great tits (*Parus major*), specifically, have become a model species for studying urban evolution across Europe and Asia and, thus, research on this species is now contributing to large collaborative research efforts that evaluate trends across replicated urban gradients in continent-wide analyses (Vaugoyeau et al. 2016; Salmón et al. 2021; Thompson et al. 2022). Despite these exciting efforts, a fundamental gap exists about whether urban phenotypic shifts in this model species are driven by evolutionary change between populations or by plastic responses to urban conditions. Common garden experiments in urban dark-eyed juncos (*Junco hymemalis*) and European blackbirds (*Turdus merula*) suggest that genetic change could at least partially play a role in phenotypic differences across morphological, physiological, and behavioural traits (Table 1; Atwell et al. 2012; Miranda et al. 2013; Reichard et al. 2020), but it remains to be seen whether this holds for other urban bird species and, specifically, the great tit where urban phenotypic shifts have been well documented in the wild.

This study uses a common garden experiment to disentangle the mechanisms that shape urban phenotypes in populations of great tits in and around Montpellier, France. In this system, we have documented several phenotypic differences between urban and forest populations in life history, morphology, physiology, and behaviour (Charmantier et al. 2017; Caizergues et al. 2018; Caizergues et al. 2022); trends that tend to be consistent across other European populations (Vaugoyeau et al. 2016; Biard et al. 2017; Corsini et al. 2021; Thompson et al. 2022). In the wild, urban great tits are smaller, have faster breath rates under constraint, show higher aggressiveness when handled, and are faster explorers than their forest counterparts, although estimates of selection gradients suggest that these urban phenotypic shifts are not favoured by natural selection (Caizergues et al. 2018; Caizergues et al. 2022). Genomic studies have revealed that, despite evident gene flow in this system, a small but significant proportion of genetic variation is explained by urbanization. This result suggests some genetic divergence between the urban and forest populations (i.e., significant F_ST_ = 0.006 – 0.009 between urban and forest comparisons; Perrier et al. 2018; Caizergues et al. 2022). A common garden approach is the next logical step in deciphering the genetic and plastic influences on the phenotypic divergences documented in this urban system.

We reared great tits from eggs collected from urban and forest sites around Montpellier under common conditions to evaluate whether documented morphological, physiological, and behavioural differences persist under the same environment. We had two major aims. The first was to compare the phenotypes of birds from urban and forest origins reared in a common garden. We hypothesized stronger genetic change for highly heritable traits (e.g., morphology, physiology) than for lowly heritable ones (e.g., behaviour; Kinnison and Hendry 2001; Stirling et al. 2002) and, therefore, that morphological and physiological differences would be more likely to persist under common conditions. More specifically, we predicted that birds from urban origins would be smaller and more stressed (phenotypic difference persists), but not more aggressive or exploratory (phenotypic difference does not persist), than birds from forest origins. Our second aim was to evaluate how different sources of variation (i.e., genetic and environmental variation) shaped phenotypes in the experimental context. We present phenotypic estimates from wild populations alongside those from the common garden for comparison.

## Methods

### Study system and quantifying urbanization

Populations of urban and forest great tits have been monitored at nest boxes in and around the city of Montpellier, France as a part of a long-term study (Charmantier et al. 2017). The forest population has been monitored since 1991 in La Rouvière forest located 20 km north of Montpellier where the number of nest boxes of 32 mm diameter entrance ranged from 37 – 119 because of theft/replacement. The urban population has been monitored since 2011 throughout the city of Montpellier at study sites that differ in their degree of urbanization (163 – 208 urban nest boxes across 8 study areas, see Figure 1 in Caizergues et al. 2024). During each spring, nest boxes are visited once per week to follow the reproduction of breeding pairs. We catch adults at nest boxes when nestlings are around 12 days old, ring them with a unique metal band, age them based on plumage as either yearling (born previous year) or adult (born at least the year before last), take a blood sample, and measure several phenotypes (see section “phenotyping”; Caizergues et al. 2018; Caizergues et al. 2022). We quantified the proportion of impervious surface area (ISA; sealed non-natural surfaces) at each nest box to generate a continuous urbanization metric to characterise the territory of breeding wild birds (captured at nest boxes) and the territory of origin for the birds raised during the common garden experiment (see supplementary methods). We also categorized territories as urban or forest habitat types since phenotypic changes may not be consistent across continuous and categorical urbanization, which can provide additional insights. For example, in cases where a phenotypic change differs between habitat types but does not clearly change along the gradient, this could suggest that i) urban effects other than ISA explain phenotypic change or ii) phenotypic changes may be non-linear.

**Figure 1.**
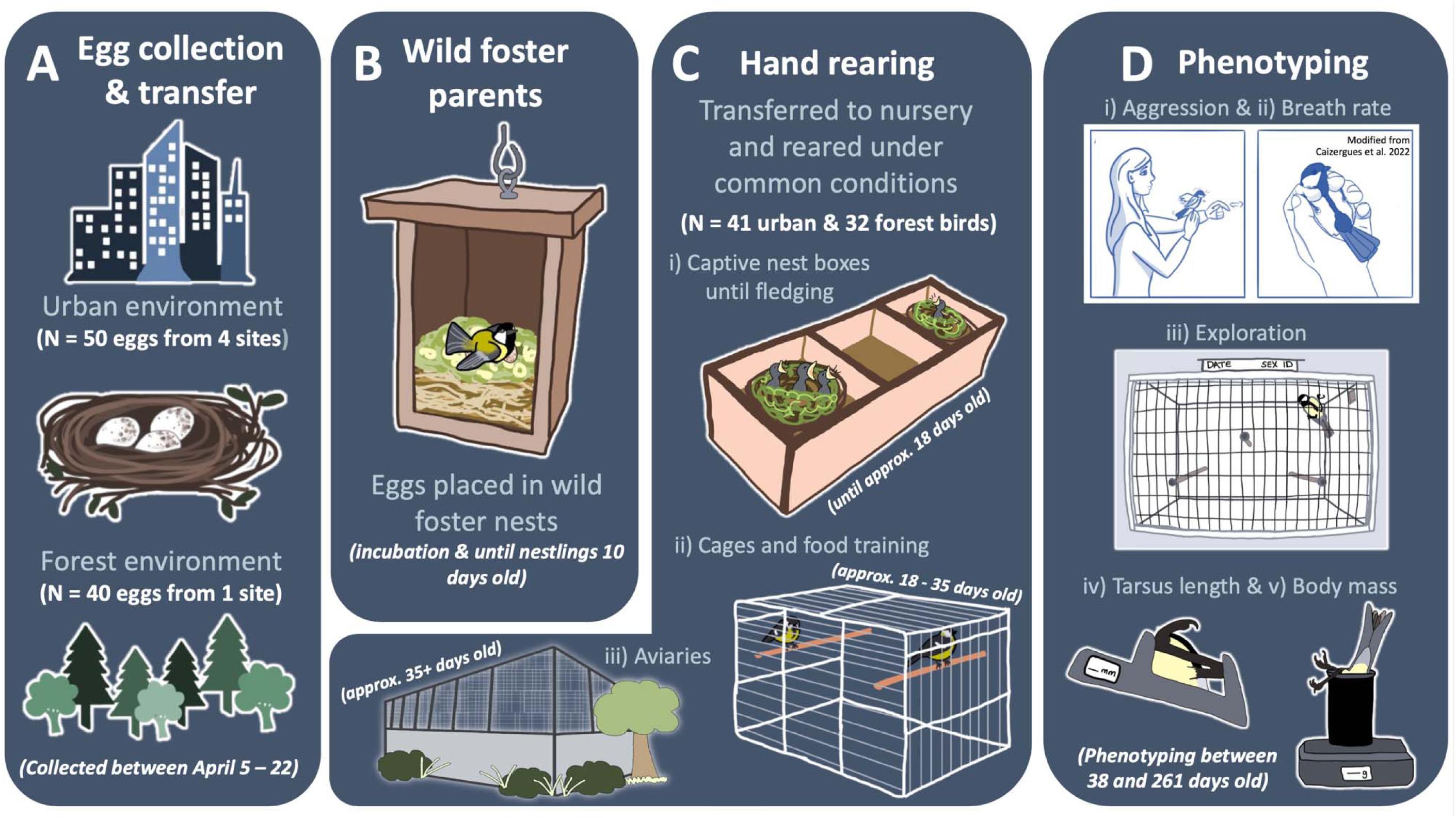
Procedure of common garden experiment including (A) egg collection, (B) transfer of eggs to wild foster parents, (C) transfer of nestlings to nursery for hand rearing under common conditions, and (D) phenotyping of common garden birds across i) handling aggression, ii) breath rate index, iii) exploration in a novel environment, iv) tarsus length, and v) body mass traits.

### Common garden manipulation

#### Egg transfer to wild foster parents

Between April 5 – 22, 2022, we collected eggs from urban (N = 50 eggs from 4 sites) and forest (N = 40 eggs from 1 site) populations (Figure 1A; Table S1). We collected three to four unincubated eggs (cold to the touch) from each origin nest box. We ensured eggs were unincubated by collecting eggs from nest boxes where we were confident that females had initiated laying within the three to four days before collection and the collected eggs were still covered by nest material. We replaced collected eggs with false eggs to encourage the origin female to continue its reproduction and we moved collected eggs into foster nest boxes at our Montpellier Zoo study site where wild females had just commenced incubation (Figure 1B). The Montpellier Zoo is an intermediate site along our urban gradient because it is natural in its vegetative characteristics, but is exposed to humans and related stimuli (Demeyrier et al. 2016). We transferred eggs from their origin to foster nests within 6 hours. In one case, we transferred eggs two days later and we kept these clutches in a dark room and rotated eggs every 12 hours until their transfer (foster ZOO46; Table S1). On average, the collected urban eggs were significantly lighter than the collected forest eggs (urban: N = 34 eggs weighed, mean = 1.56 g, variance = 0.02; forest: N = 40 eggs weighed, mean = 1.70 g, variance = 0.0075; Welch’s t-test: −4.87, df: 52.88, *P* < 0.001; Table S1).

Foster nest boxes contained eggs from two origin broods (N = 6 – 8 eggs total; Table S1). We did not mix urban and forest eggs in the same foster broods to ensure that we could confidently assign all nestlings a habitat of origin, even if biological parents abandoned or were not captured. Urban lay dates in our system were earlier than forest lay dates (urban origin nests laid on average 7.5 days earlier), but there was still overlap in reproductive phenology between habitats (Table S1). The percentage of unhatched eggs was similar across habitat of origin (18% urban and 20% forest). Unhatched forest eggs were all from one abandoned foster nest (invaded by hornets), whereas unhatched urban eggs were distributed across successful foster clutches (Table S1). Of the 90 eggs transferred, we had N = 73 nestlings hatch (41 urban and 32 forest; Table S1). We did not have mortality events so these sample sizes are representative of the number of individuals phenotyped after rearing (Table S2).

#### Captive rearing

Once nestlings could thermoregulate on their own at 10 days of age (Mertens 1977), we transferred nestlings to the Montpellier Zoo nursery between April 29 – May 16, 2022 (Figure 1C; Table S1). Due to advanced urban lay dates in our system, urban nestlings entered captivity on average 6 days before forest nestlings (urban mean = 126 Julian days, urban range = 119 – 135; forest mean = 132 Julian days, forest range = 128 – 136; Table S1). Upon arrival, we ringed and weighed nestlings before placing them into artificial nests with their foster broods. We kept them in incubators that mimicked a dim cavity and kept chicks in a quiet environment (1 – 3 broods per incubator; Figure 1C.i). At this stage, we hand-raised nestlings by feeding individuals every 30 minutes between 7:00 and 21:00 (see supplementary methods for “captive diet”). Individuals began to “fledge” their brood nests at an age of 18 days. We transferred these individuals in groups of 2-3 birds into small wire cages in the order of when they fledged (irrespective of sex, foster brood, or habitat of origin), where we trained them to feed by themselves (Figure 1C.ii). At this stage, we still fed individuals every 30 mins. Once birds were approximately 23 days old, we transferred them to larger cages (0.8 x 0.35 x 0.4 m) that allowed more movement (hops and flights) in groups of 2 individuals. At this stage their feeding schedule became less frequent, and birds were considered independent at an age of approximately 35 days when we transferred individuals to large outdoor aviaries (N = 8; size = 2.2 x 4.4 – 5.5 meters; Figure 1C.iii). Individuals were randomly organised into aviary groups blind of habitat of origin and sex (N = 6 – 10 individuals per group). All individuals were hand-reared by the same caretakers during the experiment who were blind to the origin of the birds and birds from both origins were mixed through all stages of the rearing protocol.

### Blood sampling and genotyping

We took blood samples from individuals the day before birds were transferred to outdoor aviaries to determine i) sex and nest of origin to control for genetic relatedness and ii) assign each bird with an ISA of origin. For each foster brood, nestlings had two possible nests of origin from which eggs were collected so parents of nests of origin were also blood sampled and genotyped to assign nest of origin for each common garden bird (see supplementary methods).

### Phenotypic measurements

Here we examine five phenotypic traits: handling aggression, breath rate index, exploration in a novel environment, tarsus length, and body mass. Phenotypic measurements of common garden birds (Figure 1D) followed the same protocols used to phenotype wild birds (Caizergues et al. 2018; Caizergues et al. 2022). We took all phenotypic measures of common garden birds indoors at the nursery, so all individuals were phenotyped under similar conditions (i.e., constant temperature, noise, and light levels). We took repeated measures between 06 June 2022 – 31 January 2023 and all observers were blind to habitat of origin while phenotyping individuals during four separate phenotyping sessions. On average, we measured birds 1.8 times in the wild (range = 1 – 8) and 3.4 times in the common garden (range = 1 – 4; summary by trait in supplementary and Table S2). Following our phenotyping protocol in the wild where birds are mainly measured annually during the breeding season (02 April – 16 July), we measured phenotypes at the same time in the following order:

#### Handling aggression

We measured handling aggression immediately following capture (from nest box, cage, or aviary) by provoking the bird while holding it (Figure 1D.i). We scored their aggressive response between 0 (no reaction) to 3 (tail and wings extended, pecking, and vocalization) on a scale that increased in increments of 0.5 (see Dubuc-Messier et al. 2018; Caizergues et al. 2022 for further details).

#### Breath rate index

We placed the bird in a cloth bag and allowed a five-minute standardized period of rest. Once removing the bird from the bag and properly holding the bird (Figure 1D.ii), we recorded the time it took for a bird to take 30 breaths (i.e., movements of the breast). We took this measurement twice in immediate succession and took the average between these measures to represent an individual’s breath rate index.

#### Exploration

We placed birds into a small compartment next to a novel environment arena where they had a standardized two-minute rest period. We then initiated the novel environment exploration trial by coaxing birds into the novel arena. We recorded their behaviours in this arena for 4 minutes on video then an observer later counted the number of hops and flights birds took while exploring this novel environment (Figure 1D.iii). Only one observer scored videos from the common garden (MJT) while multiple observers have scored videos from the wild populations (inter-observer reliability rho > 0.95, including MJT).

#### Tarsus length and body mass

Finally, we measured tarsus length (millimetres) with pliers to determine the length between the intertarsal notch and the end of the bent foot (i.e., Svensson’s alternative method; Svensson 1992); Figure 1D.iv) and body mass (grams) using an electronic scale (Caizergues et al. 2021; Figure 1D.v).

### Statistical analyses

We examined wild and common garden (CG) data using separate Bayesian mixed-effect models since model structures between contexts accounted for different fixed and random effects, while examining a similar main effect of interest (i.e., habitat type; aim 1). All models included habitat type (urban vs. forest) and sex (male and female), and their interaction, as fixed effects in the model to evaluate how phenotypic differences vary across habitats and sexes. If the interaction between habitat and sex was weak and largely overlapped zero, we dropped this effect and refitted the model to evaluate the phenotypic differences between habitats and sexes independently. In subsequent models, we replaced the habitat type effect with proportion ISA to further evaluate changes in the wild and CG phenotypes along a gradient of urbanization.

We have already published results on the phenotypic differences between wild urban and forest populations (Charmantier et al. 2017; Caizergues et al. 2018; Caizergues et al. 2022), but here we report estimates from wild populations that i) include more years of data (3 additional years, year range: 2011 – 2022) and ii) use data only from the study sites used in our common garden experiment (i.e., 1 forest and 4 urban study sites). All wild models included random effects that accounted for differences between individuals, study sites, and years, and we additionally accounted for differences between observers for handling aggression, breath rate index, and tarsus length (Table 2). In addition to examining habitat and sex differences in these models, we also accounted for fixed effects such as time of day, date of measurement, and protocol type following previously established model structures for these traits (Table 2; outlined by trait below; Caizergues et al. 2018; Caizergues et al. 2022).

**Table 2.**
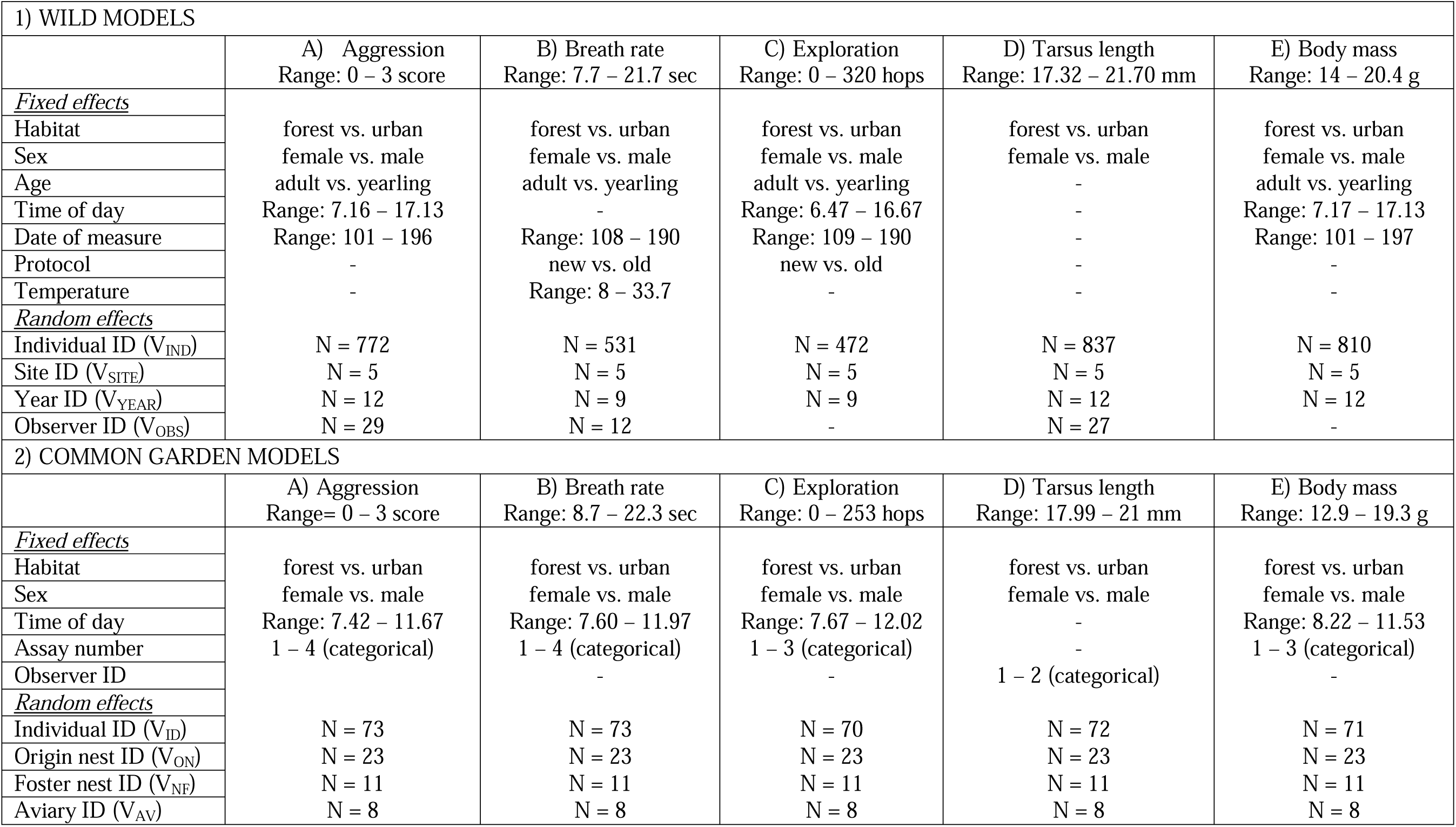
Summary of 1) wild and 2) common garden model structures that account for different fixed and random effects. Ranges for continuous fixed effects and number (N) of random effect levels for each trait are shown. Note that urbanization was also examined as a continuous effect in additional models (see model results in Table S4).

| 1) WILD MODELS |  |  |  |  |  |
| --- | --- | --- | --- | --- | --- |
|  | A) Aggression<br>Range: 0 – 3 score | B) Breath rate<br>Range: 7.7 – 21.7 sec | C) Exploration<br>Range: 0 – 320 hops | D) Tarsus length<br>Range: 17.32 – 21.70 mm | E) Body mass<br>Range: 14 – 20.4 g |
| <u>Fixed effects</u> |  |  |  |  |  |
| Habitat | forest vs. urban | forest vs. urban | forest vs. urban | forest vs. urban | forest vs. urban |
| Sex | female vs. male | female vs. male | female vs. male | female vs. male | female vs. male |
| Age | adult vs. yearling | adult vs. yearling | adult vs. yearling | - | adult vs. yearling |
| Time of day | Range: 7.16 – 17.13 | - | Range: 6.47 – 16.67 | - | Range: 7.17 – 17.13 |
| Date of measure | Range: 101 – 196 | Range: 108 – 190 | Range: 109 – 190 | - | Range: 101 – 197 |
| Protocol | - | new vs. old | new vs. old | - | - |
| Temperature | - | Range: 8 – 33.7 | - | - | - |
| <u>Random effects</u> |  |  |  |  |  |
| Individual ID (V<sub>IND</sub>) | N = 772 | N = 531 | N = 472 | N = 837 | N = 810 |
| Site ID (V<sub>SITE</sub>) | N = 5 | N = 5 | N = 5 | N = 5 | N = 5 |
| Year ID (V<sub>YEAR</sub>) | N = 12 | N = 9 | N = 9 | N = 12 | N = 12 |
| Observer ID (V<sub>OBS</sub>) | N = 29 | N = 12 | - | N = 27 | - |
| 2) COMMON GARDEN MODELS |  |  |  |  |  |
|  | A) Aggression<br>Range= 0 – 3 score | B) Breath rate<br>Range: 8.7 – 22.3 sec | C) Exploration<br>Range: 0 – 253 hops | D) Tarsus length<br>Range: 17.99 – 21 mm | E) Body mass<br>Range: 12.9 – 19.3 g |
| <u>Fixed effects</u> |  |  |  |  |  |
| Habitat | forest vs. urban | forest vs. urban | forest vs. urban | forest vs. urban | forest vs. urban |
| Sex | female vs. male | female vs. male | female vs. male | female vs. male | female vs. male |
| Time of day | Range: 7.42 – 11.67 | Range: 7.60 – 11.97 | Range: 7.67 – 12.02 | - | Range: 8.22 – 11.53 |
| Assay number | 1 – 4 (categorical) | 1 – 4 (categorical) | 1 – 3 (categorical) | - | 1 – 3 (categorical) |
| Observer ID |  | - | - | 1 – 2 (categorical) | - |
| <u>Random effects</u> |  |  |  |  |  |
| Individual ID (V<sub>ID</sub>) | N = 73 | N = 73 | N = 70 | N = 72 | N = 71 |
| Origin nest ID (V<sub>ON</sub>) | N = 23 | N = 23 | N = 23 | N = 23 | N = 23 |
| Foster nest ID (V<sub>NF</sub>) | N = 11 | N = 11 | N = 11 | N = 11 | N = 11 |
| Aviary ID (V<sub>AV</sub>) | N = 8 | N = 8 | N = 8 | N = 8 | N = 8 |

Besides examining habitat of origin effects in common garden models (aim 1), we also explored how different components of genetic and environmental variation shaped traits in the common garden (aim 2). All common garden models included the same random effects (Table 2): Individual ID accounted for variance among individuals (V_ID_), nest of origin ID accounted for variance among origin nests (V_NO_), and foster nest ID accounted for variance among foster nests (V_NF_). We also included aviary ID as a random effect that accounted for variation among social groups in the experiment (V_AV_) for the behavioural and physiological traits considered (i.e., aggression, exploration, breath rate). Since individuals in the common garden experiment were genotyped, we conducted a complementary analysis of the common garden data using mixed-effect animal models (Charmantier et al. 2014) by fitting a genetic relatedness matrix (GRM) between individuals in our common garden context (de Villemereuil et al. 2018). Since we collected unincubated eggs for the experiment, the nest of origin random effect (V_NO_) if fitted alone, may capture both genetic differences between individuals (i.e., whether they are siblings) and early environmental differences such as early maternal investment in the eggs. Therefore, the GRM approach allowed us to further evaluate how the variation of each trait was partitioned when accounting for individual genetic variation and nest of origin variation (V_A_ and V_NO_, respectively) at the same time (see supplementary methods for model description).

We computed the repeatability (*R*) of each trait in the common garden experiment as:

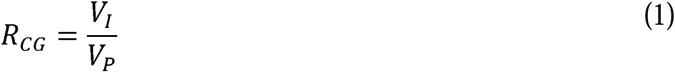

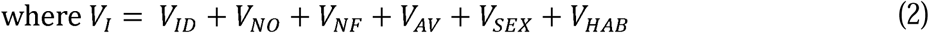

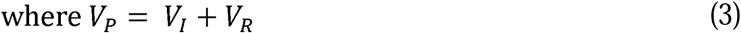

where V_I_ is the among-individual variance that comprises effects that vary consistently among individuals and drive biological differences between individuals including variance across individuals (V_ID_), nests of origin (V_NO_), foster nests (V_NF_), aviaries or social groups (V_AV_), and fixed effect variance among sexes and habitat types (V_SEX_ and V_HAB_). We only chose to include fixed effect variance generated by biological effects in the model (i.e., habitat and sex), rather than experimental effects (e.g., time of day or observer), to quantify repeatability using natural sources of variation that may improve comparability to other studies (de Villemereuil, Morrissey, et al. 2018; Wilson 2018). V_P_ is the total phenotypic variance and includes sources of among-individual variance (V_I_) and residual variance (V_R_). For comparison, we computed *R* in wild birds as:

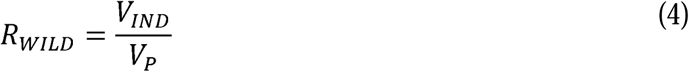

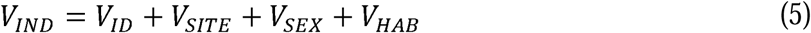

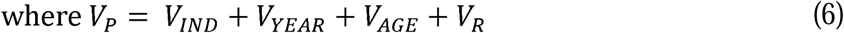

where V_IND_ is the among-individual variance and comprises variance across individuals (V_ID_), study sites or local habitats (V_SITE_), and sexes and habitat types (fixed effect variance: V_SEX_ and V_HAB_). V_P_ is the total phenotypic variance and includes among-individual variance (V_IND_) and residual variance (V_R_). In the wild context, we also incorporated biological effects in V_P_ that vary within individuals including variance among years (V_YEAR_) and fixed effect variance among ages (V_AGE_). For Poisson models (i.e., exploration) that use a log link transformation, we used the QCglmm package (de Villemereuil et al. 2016) to convert the variance components and repeatability estimate from the latent scale to the data scale.

We conducted all analyses in R v4.3.3 (R Core Team 2024) using Bayesian mixed-effect models in the MCMCglmm package (Hadfield 2010) using a Gaussian error structure for all traits except exploration (number of hops) where we used a Poisson error structure. See supplementary methods and Table 2 for description of other fixed effects included for each model. We used weakly informative inverse-Gamma priors (V = 1, nu = 0.002) for fixed and random effects. We ran all models for 1000000 iterations, with a thinning of 500 and a burn-in period of 10000, which achieved effective sample sizes > 1000 across all estimates. We verified model fit by visually inspecting histograms and QQPlots of model residuals, and the relationship between the residuals and fitted values. We confirmed convergence of models by visually inspecting trace plots, verifying low autocorrelation, and by using Heidelberg stationary tests (de Villemereuil 2018).

## Results

### Aggression in hand

We found clear evidence that wild urban males were more aggressive than wild forest males, but no habitat difference for females (habitat*sex effect, Table 3.1A; Figure 2A). Results in the wild were qualitatively similar when examining how phenotypes changed along the urban gradient; there was clear evidence that wild males in habitats with higher proportion ISA had increased aggression (Table S4; Figure 3A). There was no clear evidence that this phenotypic difference was maintained in the common garden (weak habitat*sex or habitat effects overlapping zero; Table 3.2A; Table S5; Figure 2A), and no clear evidence that handling aggression increased with proportion ISA of the origin habitat of common garden birds (credible interval overlaps zero; Table S4). Individual ID explained 23% of the variation in aggression in the common garden, nest of origin explained 2%, foster nest explained 2%, and housing aviary explained 2% of the variation (Table 3.2A).

**Figure 2.**
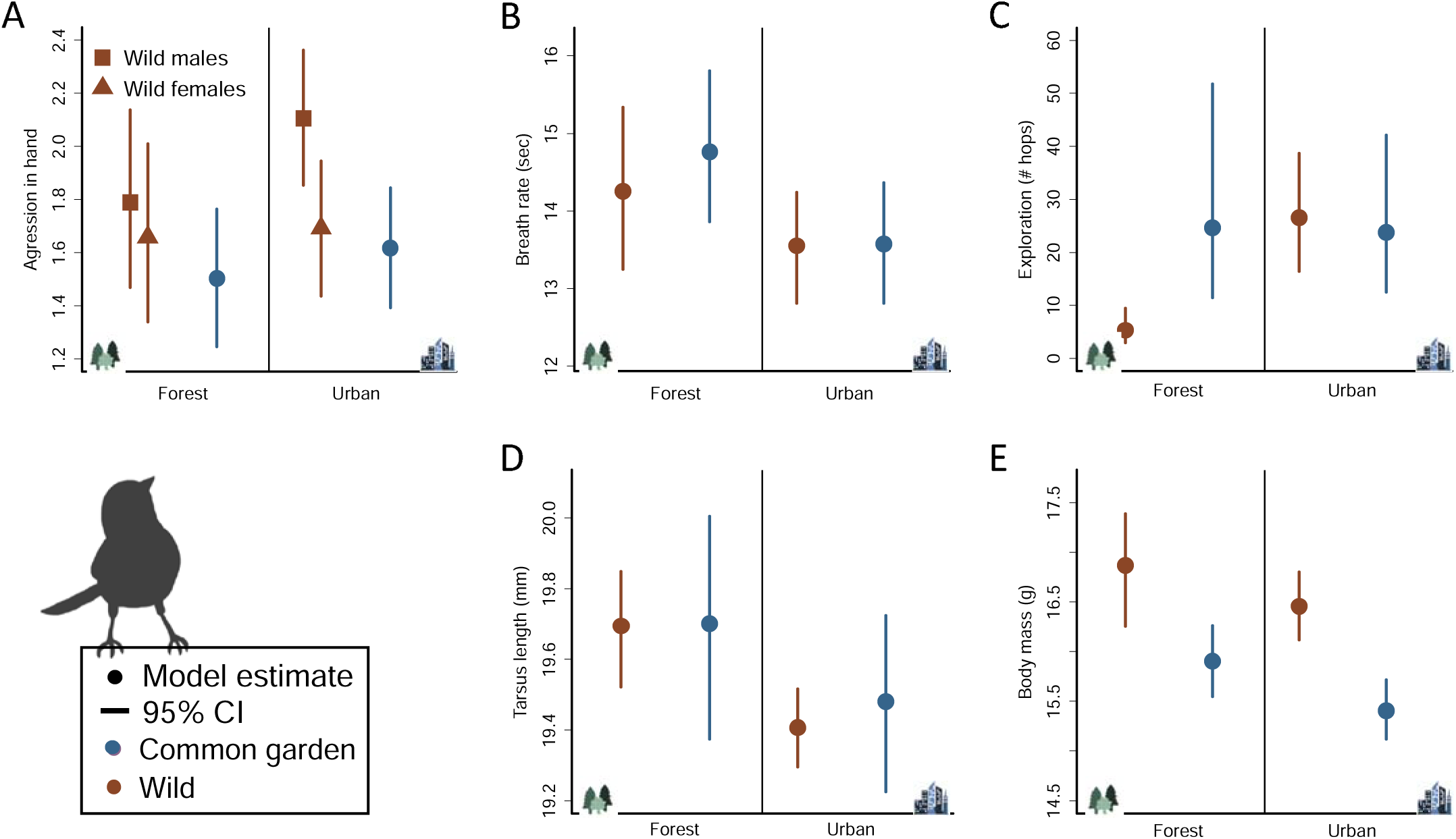
Habitat type model estimate and 95% credible intervals (CI) on phenotypic traits of wild (brown) and common garden birds (blue) across A) aggression in hand, B) breath rate index, C) exploration score, D) tarsus length, and E) body mass. Habitat differences varied clearly by sex only in one case (A: aggression in wild birds) and these sex differences are shown (wild males: squares, wild females: triangles); aggression did not differ clearly by sex in the common garden. Phenotypes were measured in the wild annually during the breeding season between 2011– 2022, whereas we measured phenotypes in the common garden between 06 June 2022 – 31 January 2023. See also Figure S1 for plots that show raw data.

**Figure 3.**
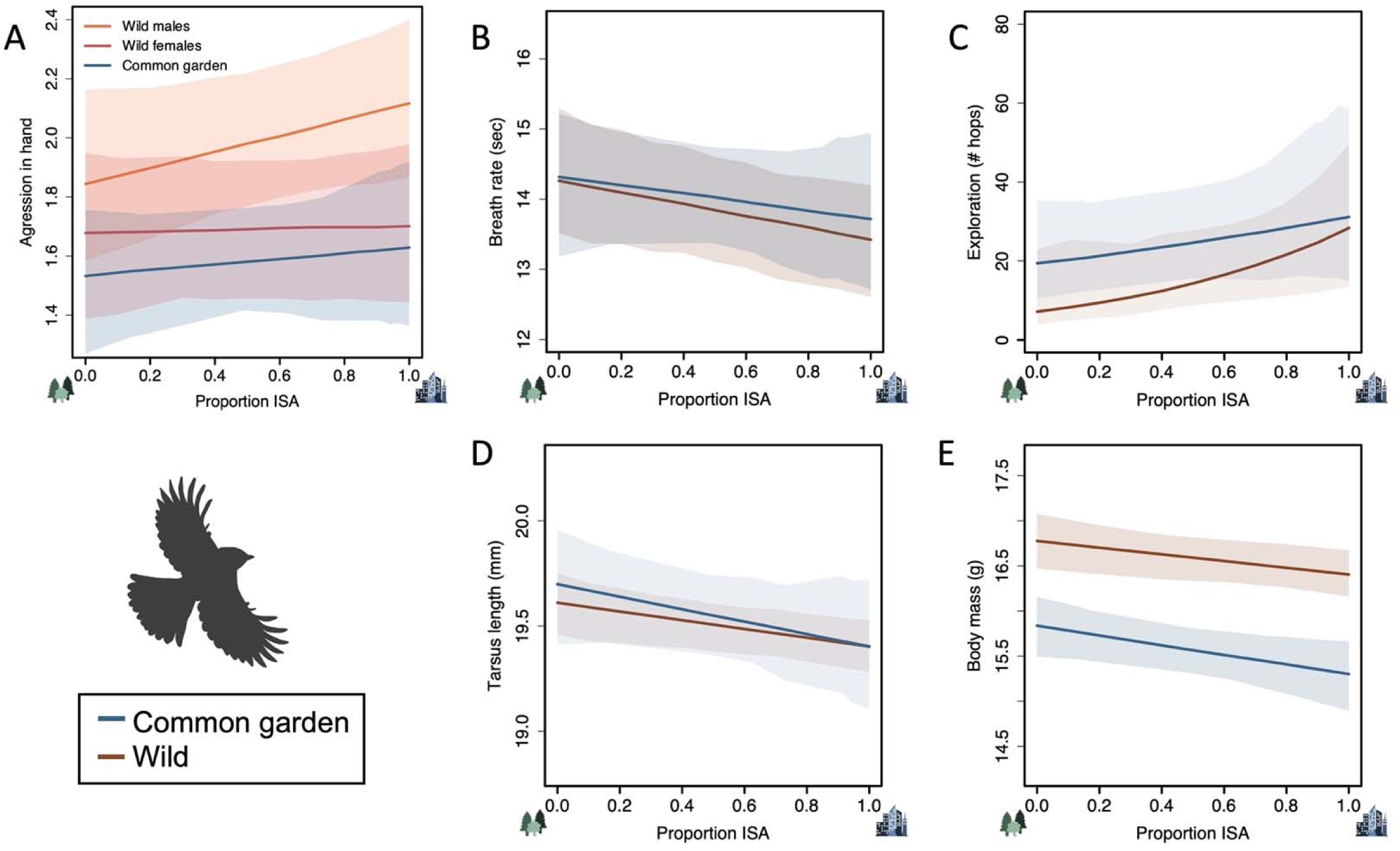
Proportion of impervious surface area (ISA) model estimate and 95% credible intervals (CI) on phenotypic traits of wild (brown) and common garden (blue) birds across A) aggression in hand, B) breath rate index, C) exploration score, D) tarsus length, and E) body mass. ISA effects varied clearly by sex only in one case (A: aggression in wild birds) and these sex differences are shown (wild males: orange, wild females: red); aggression over the ISA gradient in the common garden did not differ clearly by sex. Phenotypes were measured in the wild annually during the breeding season between 2011– 2022, whereas we measured phenotypes in the common garden between 06 June 2022 – 31 January 2023. See also Figure S2 for plots that show raw data.

**Table 3.** Fixed and random effect estimates and credible intervals (CI) for 1) wild and 2) common garden contexts across phenotypic traits (A: handling aggression, B: breath rate index, C: exploration, D: tarsus length, and E: body mass). Exploration estimates are from a Poisson generalized mixed-effect model, while all other traits were fit with Gaussian mixed-effect models. Common garden models estimated Individual ID (V_ID_), origin nest ID (V_NO_), foster nest ID (V_NF_), aviary ID (V_AV_), and residual variance (V_R_). The number of observations (obs) and individuals (ind) for each trait and context are shown. Shown in bold are fixed effects whose credible intervals exclude zero or random effects whose lower CI is ≥0.001. Computed among-individual variance (V_IND_= V_ID_+V_SITE_+V_SEX_+V_HAB_, V_ID_=V_ID_+V_NO_+V_NF_+V_AV_+V_SEX_+V_HAB_; equations 2&5) and repeatability (*R*) are shown for both contexts for comparison.

| 1) WILD |  |  |  |  |  |  |  |  |  |  |
| --- | --- | --- | --- | --- | --- | --- | --- | --- | --- | --- |
|  | A) Aggression<br>N = 1308 obs, 772 ind |  | B) Breath rate<br>N = 702 obs, 531 ind |  | C) Exploration<br>N = 581 obs, 472 ind |  | D) Tarsus length<br>N = 1437 obs, 837 ind |  | E) Body mass<br>N = 1375 obs, 810 ind |  |
| *Fixed effects* | Est | CI | Est | CI | Est | CI | Est | CI | Est | CI |
| Intercept | 2.29 | 1.68 - 2.84 | 12.74 | 10.53 - 15.19 | 3.32 | 0.78 - 5.97 | 19.42 | 19.25 - 19.58 | 15.99 | 15.35 - 16.65 |
| Habitat (urban) | 0.03 | -0.3 - 0.46 | -0.70 | -1.71 - 0.35 | **1.60** | **0.84 - 2.2** | **-0.29** | **-0.48 - -0.11** | -0.41 | -1.14 - 0.18 |
| Sex (male) | 0.13 | -0.04 - 0.3 | 0.06 | -0.3 - 0.4 | -0.06 | -0.47 - 0.32 | **0.54** | **0.48 - 0.62** | **0.60** | **0.5 - 0.7** |
| Age (yearling) | -0.09 | -0.2 - 0.02 | -0.008 | -0.32 - 0.29 | 0.13 | -0.19 - 0.48 |  |  | **-0.31** | **-0.39 - -0.24** |
| Time of day | **-0.04** | **-0.06 - -0.01** |  |  | -0.01 | -0.1 - 0.07 |  |  | **0.04** | **0.02 - 0.06** |
| Date of measure | -0.001 | -0.003 - 0.001 | -0.003 | -0.02 - 0.01 | -0.01 | -0.03 - 0.01 |  |  | 0.003 | 0.001 – 0.005 |
| Protocol (old) |  |  | 0.16 | -0.39 - 0.71 | 0.16 | -0.35 - 0.63 |  |  |  |  |
| Temperature |  |  | **0.09** | **0.06 - 0.13** |  |  |  |  |  |  |
| Habitat * Sex | **0.28** | **0.03 - 0.53** |  |  |  |  |  |  |  |  |
| *Random effects* |  |  |  |  |  |  |  |  |  |  |
| Individual ID (V<sub>IND</sub>) | **0.43** | **0.34 - 0.52** | **2.61** | **2.06 - 3.19** | **2.73** | **1.99 - 3.5** | **0.28** | **0.25 - 0.31** | **0.39** | **0.33 - 0.45** |
| Site ID (V<sub>SITE</sub>) | 0.02 | <0.001 - 0.08 | 0.25 | <0.001 - 0.74 | 0.08 | <0.001 - 0.3 | 0.008 | <0.001 - 0.02 | 0.09 | 0.001 - 0.3 |
| Year ID (V<sub>YEAR</sub>) | 0.03 | <0.001 - 0.07 | 0.05 | <0.001 - 0.19 | 0.06 | <0.001 - 0.22 | 0.002 | <0.001 - 0.004 | **0.06** | **0.01 - 0.13** |
| Observer ID (V<sub>OBS</sub>) | **0.04** | **0.01 - 0.08** | **0.52** | **0.1 - 1.26** |  |  | 0.005 | 0.001 - 0.01 |  |  |
| Residual variance (V<sub>R</sub>) | **0.55** | **0.48 - 0.62** | **1.75** | **1.4 - 2.1** | **1.61** | **1.08 - 2.14** | **0.02** | **0.02 - 0.02** | **0.27** | **0.24 - 0.31** |
| V<sub>IND</sub> | **0.48** | **0.38 – 0.70** | **2.89** | **2.28 – 4.36** | **3.44** | **2.57 – 4.61** | **0.38** | **0.34 – 0.44** | **0.57** | **0.47 – 1.15** |
| R<sub>WILD</sub> | **0.45** | **0.37 – 0.56** | **0.62** | **0.52 – 0.73** | **0.13** | **0.06 – 0.20** | **0.94** | **0.93 – 0.95** | **0.62** | **0.53 – 0.78** |
| 2) COMMON GARDEN |  |  |  |  |  |  |  |  |  |  |
|  | A) Aggression<br>N = 285 obs, 73 ind |  | B) Breath rate<br>N = 283 obs, 73 ind |  | C) Exploration<br>N = 203 obs, 70 ind |  | D) Tarsus length<br>N = 211 obs, 72 ind |  | E) Body mass<br>N = 210 obs, 71 ind |  |
| *Fixed effects* | Est | CI | Est | CI | Est | CI | Est | CI | Est | CI |
| Intercept | 1.40 | 0.23 - 2.59 | 9.43 | 7.21 - 11.62 | 4.02 | 1.91 - 6.07 | 19.43 | 19.1-19.76 | 15.84 | 14.71 - 16.98 |
| Habitat (urban) | 0.12 | -0.2 - 0.45 | **-1.16** | **-2.32 - -0.02** | -0.05 | -1.06 - 0.81 | -0.22 | -0.64 - 0.17 | **-0.50** | **-0.96 - -0.03** |
| Sex (male) | -0.23 | -0.5 - 0.05 | **0.81** | **0.03 - 1.6** | -0.36 | -0.84 - 0.14 | **0.46** | **0.24 - 0.67** | **1.03** | **0.75 - 1.33** |
| Time of day | 0.04 | -0.07 - 0.16 | **0.47** | **0.27 - 0.67** | -0.05 | -0.24 - 0.12 |  |  | -0.10 | -0.2 - 0.02 |
| Measurement (2) | -0.17 | -0.44 - 0.12 | **0.94** | **0.5 - 1.37** | -0.18 | -0.67 - 0.24 |  |  | 0.12 | -0.04 - 0.28 |
| Measurement (3) | -0.25 | -0.52 - 0.01 | **0.50** | **0.04 - 0.9** | -0.13 | -0.58 - 0.3 |  |  | **0.50** | **0.37 - 0.67** |
| Measurement (4) | -0.38 | -0.62 - -0.12 | -0.38 | -0.8 - 0.02 |  |  |  |  |  |  |
| Observer (2) |  |  |  |  |  |  | **0.07** | **0.04 - 0.11** |  |  |
| *Random effects* |  |  |  |  |  |  |  |  |  |  |
| Individual ID (V<sub>ID</sub>) | **0.19** | **0.08 - 0.32** | **2.15** | **1.1 - 3.18** | **0.47** | **<0.001 - 0.98** | **0.18** | **0.11 - 0.27** | **0.26** | **0.13 - 0.41** |
| Origin nest ID (V<sub>NO</sub>) | 0.02 | <0.001 - 0.06 | 0.58 | <0.001 - 1.66 | 0.20 | <0.001 - 0.71 | 0.10 | <0.001 - 0.22 | 0.12 | <0.001 - 0.3 |
| Foster nest ID (V<sub>NF</sub>) | 0.01 | <0.001 - 0.05 | 0.15 | <0.001 - 0.68 | 0.28 | <0.001 - 1.1 | 0.03 | <0.001 - 0.1 | 0.04 | <0.001 - 0.13 |
| Aviary ID (V<sub>AV</sub>) | 0.01 | <0.001 - 0.05 | 0.14 | <0.001 - 0.62 | 0.05 | <0.001 - 0.21 |  |  |  |  |
| Residual variance (V<sub>R</sub>) | **0.56** | **0.46 - 0.67** | **1.58** | **1.29 - 1.87** | **1.66** | **1.22 - 2.17** | **0.01** | **0.01-0.02** | **0.18** | **0.14 - 0.23** |
| V<sub>I</sub> | **0.25** | **0.14 – 0.45** | **3.74** | **2.54 – 5.83** | **1.01** | **0.39 – 2.49** | **0.38** | **0.26 – 0.60** | **0.74** | **0.53 – 1.11** |
| R<sub>CG</sub> | **0.30** | **0.17 – 0.44** | **0.68** | **0.54 – 0.78** | **0.10** | **0.03 – 0.17** | **0.97** | **0.95 – 0.98** | **0.80** | **0.73– 0.87** |

### Breath rate index

We found weak evidence that wild urban birds had faster breath rates than wild forest birds (6% posterior distribution crossing zero, Table 3.1B; Figure 2B), but clear evidence that the amount of time to take 30 breaths decreased across the urbanization gradient in the wild populations (i.e., faster breath rates in more urbanized habitats, credible interval excludes zero, Table S4; Figure 3B). We found clear evidence that this phenotypic difference was maintained in common garden birds where birds from urban origins had faster breath rates than birds from forest origins (Table 3.2B; Figure 2B), but there was no clear change in breath rate across the urbanization gradient in the common garden (credible interval overlaps zero; Table S4; Figure 3B). Individual ID explained 40% of the variation in breath rates in the common garden, while origin nest (11%), foster nest (4%), and housing aviary (4%) explained less variation in this trait (Table 3.2B).

### Exploration

We found clear evidence that exploration in the wild was higher in urban compared to forest birds (Table 3.1C; Figure 2C) and less clear evidence that exploration increased with increasing urbanization (5% posterior distribution crossing zero; Table S4; Figure 3C). There was no clear evidence that these differences were maintained in the common garden (credible overlapping zero; Table 3.2C; Figure 2C), and exploration of common garden birds did not increase clearly with proportion ISA of the origin habitat (Table S4; Figure 3C). Individual ID explained 17% of the variation in common garden exploration behaviours while origin nest (7%), foster nest (10%), and housing aviary (2%) explained less variation in this trait (Table 3.2C).

### Tarsus length

We found clear evidence that tarsus length was significantly shorter in wild urban birds than wild forest conspecifics (Table 3.1D; Figure 2D) and with increasing urbanization in the wild (Table S4; Figure 3D). In the common garden experiment, this phenotypic divergence was in the same direction and had similar effect sizes as in the wild, with birds from urban origins tending to have shorter tarsi than birds from forest origins (CG: β_habitat_ = −0.22 vs. wild: β_habitat_ = −0.29; Table 3D) and tarsus length decreasing with increasing ISA (CG: β_ISA_ = −0.29 vs. wild: β_ISA_ = −0.20, Table S4). The large credible intervals in the common garden context however reduced certainty and suggests the smaller common garden sample size may limit the ability to detect statistical differences for this trait (CI overlapping zero, but note 91% of ISA CI excludes 0). Out of the random effects considered, individual ID explained 46% of the variation in tarsus in the common garden, followed by origin nest and foster nest which explained 25% and 7% of the variation, respectively (Table 3.2D).

### Body mass

We found weak evidence that wild urban birds were lighter than wild forest birds (7% posterior crossing zero; Table 3.1E; Figure 2E), but clear evidence that body mass decreased with increasing urbanization (credible interval excluded zero, Table S4; Figure 3E). We found clear evidence that common garden birds from urban origins were lighter than birds from forest origins (Table 3.2E; Figure 2E). Results in the common garden across the urban gradient were consistent with this conclusion where the weight of common garden birds decreased with increasing urbanization of the origin habitat (Table S4; Figure 3E). Individual ID explained 28% variation in common garden body mass, followed by origin nest which explained 12% variation, and foster nest effects which explained 4% (Table 3.2E). On average common garden birds were significantly lighter than wild birds, reflecting an experimental effect on body mass (Welch’s t-test: t = −22.30, df = 272, *P* < 0.001, Figure 2E; Figure 3E).

### Additive genetic variance and heritability of common garden traits

The complementary quantitative genetic models (Table S3) revealed that breath rate index, tarsus length, and body mass measured in common garden birds were moderately heritable (breath rate: V_A_ = 1.53, *h^2^* = 0.23 [0.001 – 0.57]; tarsus length: V_A_ = 0.16, *h^2^* = 0.33 [0.003 – 0.72]; body mass: V_A_ = 0.28, *h^2^* = 0.28 [0.01 – 0.49]), while aggression in hand and exploration were lowly heritable (aggression: V_A_ = 0.07, *h^2^* = 0.05 [<0.001– 0.28]; exploration: V_A_ = 0.43, *h^2^* = 0.02 [<0.001 – 0.06]). Although there was uncertainty around variance estimates in these models (i.e., credible intervals close to zero), estimates of V_NO_ were low across all traits (Table S3) providing additional insight into the minimal contributions from early maternal effects in the experiment.

## Discussion

We found evidence that both genetic and plastic changes have contributed to phenotypic shifts in wild urban tits, but the relative contributions of these drivers depend on the trait (aim 1). Specifically, our results provide evidence that genetic differences between populations have strongly driven the divergence observed in breath rate and body mass, while plasticity to urban conditions predominately contributes to divergences in aggression and exploration. Further, we found that individual differences tended to explain the most trait variation in the experiment, whereas nest of origin and foster nest variation had minimal contributions (aim 2).

We found support that genetic change or very early maternal investment in eggs has driven population divergences in breath rate index as phenotypic differences between birds from urban and forest origins were clearly maintained in our experiment (but note difference not statistically supported along gradient). In line with findings in the wild populations (Caizergues et al. 2022), birds from urban origins had faster breath rates than birds from forest origins. As breath rate index correlates with heart rate under constraint (Dubuc-Messier et al. 2016) and has previously been associated with physiological stress responses in this species (Carere and van Oers 2004; Krams et al. 2014), our results could indicate that genetic change in urban populations has contributed to a more proactive coping style in urban environments (Koolhaas et al. 2010; Koolhaas et al. 2011). Our results differ from those in urban European blackbirds (*Turdus merula*; Partecke et al. 2006) and juncos (*Junco hyemalis*; Atwell et al. 2012) where lower stress responses of urban birds were maintained when individuals were reared under common conditions. Indeed, there is no general consensus on how urbanization impacts stress responses in birds (reviewed in Bonier 2012) and so our results make a useful contribution towards understanding how physiology might impact adaptation to urban contexts.

We also found support that being smaller may have a genetic basis in cities. Specifically, birds from urban origins were lighter than birds from forest origins, despite being fed the same diet *ad libitum*. This habitat difference in body mass was statistically clear and higher in the common garden than the wild supporting a genetic basis for shifts to smaller urban body mass, rather than plasticity which could possibly reduce this wild phenotypic difference. Lighter urban body mass in the experiment could also be explained by early maternal investment in the egg, especially since the urban eggs collected for the experiment were on average lighter than forest eggs. Since egg size is highly heritable (e.g., egg volume h^2^ = 0.6 – 0.8, Van Noordwijk et al. 1981; Hõrak et al. 1995; see also Christians 2002) and female body size can positively correlate with their egg size in this species (Hõrak et al. 1995) it is possible that genetic differences between females in maternal egg investment could shape body mass variation in the wild.

Birds from urban origins also tended to have smaller tarsi than birds from forest origins in line with the phenotypic shift from the wild (Caizergues et al. 2018) but this was not supported statistically. We could lack statistical power to make firm conclusions on whether this difference in tarsus length was maintained in the common garden, especially since the wild phenotypic difference in tarsus length is small and would require large samples to detect. The habitat difference for tarsus length was weaker in the common garden than the wild (i.e., difference of 0.22 mm in common garden vs 0.29 mm in wild) which may indicate that a combination of plastic and genetic effects explain the tarsus length difference. Tarsus length and body mass are heritable traits and tend to strongly correlate with each other in this species (Gebhardt-Henrich and Van Noordwijk 1991; Hõrak 1994; Gosler and Harper 2000; Young and Postma 2023), suggesting that parallel genetic change for tarsus alongside body mass is possible. However, tarsus development is also strongly influenced by environmental conditions in early life (Dhondt 1982; Merilä and Fry 1998; Talloen et al. 2010; Seress et al. 2020), and so we hypothesize that genetic and plastic effects both contribute to smaller tarsus lengths in urban birds. Further quantitative genetic approaches using long-term datasets on wild populations, and observed or genetically reconstructed pedigrees, will provide a useful complementary exploration on the underlying drivers behind shifts to smaller urban tarsus lengths in tits.

Decreases in traits associated with body size are documented across diverse taxa in cities (Merckx et al. 2018; Hahs et al. 2023) and this phenotypic shift is hypothesized to facilitate heat dissipation and be an adaptive response to rising global temperatures that are pronounced in urban areas via the heat island effect (Youngflesh et al. 2022; Sumasgutner et al. 2023). For example, *Daphnia* from urban origins had smaller body sizes and higher heat tolerance in a common garden experiment than those from nonurban origins, and there was evidence that smaller urban body sizes could indirectly increase heat tolerance in this species (Brans et al. 2017). Further, city great tits tend to be lighter than forest tits across Europe (Thompson et al. 2022) and, in Veszprém, reproduction of city tits is less affected by extreme temperature than their forest counterparts (Pipoly et al. 2022). These results suggest that city great tits could be adapted to warming conditions and our results could imply that decreases in urban body size are an adaptive response to heat island effects (but see Playà Montmany et al. 2021). Smaller body size does not appear to afford urban great tits in Montpellier reproductive benefits (Caizergues et al. 2018), hence further work will be needed to evaluate whether smaller body sizes, or other correlated traits, are associated with higher survival in urban habitats.

Genetic change between populations that contribute to phenotypic differences can also arise via neutral evolutionary processes like genetic drift or founder effects (Leinonen et al. 2013), and differentiating these processes from local adaptation is informative to evaluate whether populations are adapting in pace with environmental change (de Villemereuil et al. 2020). We found evidence that breath rate and body mass differences are likely driven by genetic change or very early maternal investment, but we are unable to completely dismiss the role of neutral evolutionary processes towards genetic differences between populations. Using a complementary quantitative genetics approach, we estimated higher genetic differences underlying these traits (computed Q_ST_ values in Table S3: 0.06 and 0.08) than would be expected by neutral genetic variation between these populations (F_ST_ values between 0.006 – 0.008; Perrier et al. 2018). However, the uncertainty around these Q_ST_ estimates (credible intervals crossed 0.006) prevent us from completely excluding neutral evolutionary processes here. In future, rearing individuals from multiple city and forest comparisons in a common garden experiment would further strengthen our evidence against processes of neutral evolution and possibly demonstrate parallel evolutionary trajectories across multiple city populations.

We did not find evidence that genetic change has considerably contributed to urban behavioural shifts as birds from urban and forest origins did not clearly differ in their aggressive or exploratory behaviours in the experiment. It is commonly assumed that urban populations are evolving and adapting to novel urban conditions (Lambert et al. 2021), and behavioural adaptations may be particularly important in this process (Miranda et al. 2013). Alternatively, it has been argued that phenotypic adjustments through plasticity are probably more frequent (Hendry et al. 2008), especially for behavioural traits (Sol et al. 2013; Caspi et al. 2022). Our results provide support for the latter argument and contrast findings in urban blackbirds and juncos where behavioural differences were assumed to be a result of local adaptation (Atwell et al. 2012; Miranda et al. 2013). We therefore conclude that more aggressive and exploratory behaviours of wild urban great tits are most strongly driven by plastic adjustments to life in cities. Habitat matching behaviours (Edelaar et al. 2008; Camacho et al. 2020) could contribute to these urban differences in the wild, whereby more aggressive and exploratory individuals disperse and settle in more urbanized habitats, if these behaviours provide them an advantage in urban habitats. However, these behaviors were not found to covary at the individual level in an urban behavioral syndrome and do not seem to afford reproductive or survival benefits in the urban population (Caizergues, Grégoire, et al. 2022). Dispersal dynamics and habitat matching behaviours in an urban context are still poorly understood but, as these behaviors do not covary or improve fitness, we so far have limited evidence that habitat matching contributes to these urban phenotypic shifts.

Our second aim was to investigate how different sources of environmental and genetic variation contributed to repeatable individual differences across traits in our experimental context. The estimated among-individual variance and repeatability of traits in the common garden were remarkably similar to those estimated in the wild (i.e., similar estimates and overlapping credible intervals). Trait variation in the experiment tended to be shaped predominately by differences between individuals (i.e., Individual ID). Individual ID in the experiment could comprise both individual-specific genetic and environmentally induced individual differences, and our complementary quantitative genetic analysis suggested that individual genetic variation explains between 23 – 33% of the variation across breath rate, tarsus length and body mass, but only 2 – 5% for aggression and exploration behaviours (i.e., estimates of V_A_ in Table S3, but note wide credible intervals). Variation attributed to foster nest and aviary, likely related to brood and social environmental conditions respectively, remained low across traits (3 – 14%). Specifically, early environmental conditions can affect tarsus development and growth (e.g., Seress et al. 2020), but we found limited support that foster parents and nests contributed to individual differences in tarsus. Overall, estimated individual differences were similar between common garden and wild contexts, especially in those traits where we find evidence for underlying genetic differences between populations.

Finally, a few caveats should be considered when interpreting our results. First, we are unable to fully discount the contribution of very early maternal effects towards the maintained breath rate and body mass differences in the common garden. By collecting unincubated eggs we limited maternal contributions to egg investment, which could influence morphological traits like body mass (Hõrak et al. 1995). Although we found limited (but unclear) support for maternal effects in our experiment (i.e., negligible V_NO_; Table S3), our results should be interpreted with this in mind. Second, birds in our experiment were assayed at a relatively young age (between 38 - 261 days old) compared to when they are usually assayed in the wild (73% observations at 1 year old), which may affect how our common garden estimates compare to our wild populations. However, measuring phenotypic traits earlier in our experiment seemed to have limited impact on results as most common garden phenotypes were similar to wild phenotypes. Body mass in the common garden was the only trait that seemed to differ from the wild. Wild juvenile birds (1 year of age) tend to be on average 0.3 g lighter than wild adults in this population, so this may indicate that age could at least partially contribute to the observed difference in mass between contexts. Third, we monitor forest great tits in one larger study site and, although this nonurban area contains different forest types, we lack replication to draw broad inferences on the phenotypes of forest great tits more generally. Fourth, although the effects of urbanization on wild trait divergences were in the same direction and of similar magnitude to what has been previously reported in these populations, there were three cases in this study where statistical support varied (statistically unclear effect of habitat on breath rate and body mass, and ISA on exploration). In all these cases >93% of the posterior was negative (positive for exploration) indicating weak tendencies, which were likely driven by methodological differences between this and previous studies (i.e., different subset of data, additional years of data, Bayesian approach). Finally, rearing individuals under the same restricted and benign conditions (e.g. *ad libitum* food supply) may have prevented us from detecting phenotypic differences if they are impacted by genetic and environmental interactions (G x E; Conover et al. 2009). Although difficult to conduct, multi-treatment common gardens where food or temperature are manipulated could be especially valuable for teasing apart genetic and environmental interactions acting on phenotypic shifts in urban great tits.

In conclusion, our survey of the literature for urban common garden experiments indicates that both plastic and genetic divergences between urban and nonurban populations are common. In our study we find evidence that urban phenotypic divergences in stress physiology and morphology are mainly driven by genetic change or very early maternal investment in eggs. Common gardens are not able to affirm local adaptation, unless realistic multi-treatment or reciprocal transplant approaches are used (e.g., Gorton et al. 2018; Tüzün and Stoks 2021), and evaluating reproductive and survival benefits of our common garden birds in aviaries would not be appropriate. Thus, investigating whether these genetic differences between populations are adaptive remains an avenue for future research. We did not find evidence that genetic change strongly drives urban behavioural shifts, which provides contrary evidence to urban common garden studies in other bird species (Atwell et al. 2012; Miranda et al. 2013). Further work will be needed to uncover whether plasticity predominantly drives other urban behaviours in great tits (e.g., neophilia or boldness) and determine the mechanisms underlying discrepancies with other studies. Our results highlight that phenotypic shifts in urban populations can be impacted by both genetic and plastic changes and make a valuable contribution in filling a fundamental gap concerning the urban evolution of a model species. Examining how evolutionary processes in urban contexts impact phenotypic and genetic variation will have important applications for conserving urban wildlife populations and their ecological roles in communities (Lambert and Donihue 2020; Des Roches et al. 2021), but will also improve our fundamental understanding of ecology and evolution in wild systems more broadly, especially in light of global environmental change.

## Acknowledgements

We would like to thank everyone who participated in the common garden and is not an author: Laura Gervais, Barbara Class, Dhanya Bharath, Christophe de Franceschi, Sam Perret from the CEFE and Vivian Espinasse, Thibault Pujol, Flavien Daunis, Cathie Troussier, Jérôme Brière, Laetitia Boscardin, Sébastien Pouvreau, Lucas Boussioux, Charlotte Gay, Marion Darde from the Montpellier Zoo. Thanks to Pierre de Villemereuil for useful discussions and coding help. Comments and suggestions on a previous version from Timothée Bonnet and an anonymous reviewer improved the content and clarity of the manuscript.

## Author contributions

MJT, DR, SPC, and AC conceived the study. All authors developed the study methodology and collected data. MJT compiled the data, performed the analyses, and wrote the first draft of the manuscript. MJT, DR, SPC, and AC interpreted the results. All authors contributed substantially to reviewing and editing drafts of the manuscript.

## References

Alberti M, Marzluff J, Hunt VM. 2017. Urban driven phenotypic changes: empirical observations and theoretical implications for eco-evolutionary feedback. Philosophical Transactions of the Royal Society B: Biological Sciences. 372(1712):20160029. doi:10.1098/rstb.2016.0029.

Altermatt F, Ebert D. 2016. Reduced flight-to-light behaviour of moth populations exposed to long-term urban light pollution. Biology letters. 12(4):20160111. 10.1098/rsbl.2016.0111.

Antonio-Nkondjio C, Fossog BT, Ndo C, Djantio BM, Togouet SZ, Awono-Ambene P, Costantini C, Wondji CS, Ranson H. 2011. *Anopheles gambiae* distribution and insecticide resistance in the cities of Douala and Yaoundé (Cameroon): influence of urban agriculture and pollution. Malaria Journal. 10(1):1–13.

Atwell JW, Cardoso GC, Whittaker DJ, Campbell-Nelson S, Robertson KW, Ketterson ED. 2012. Boldness behavior and stress physiology in a novel urban environment suggest rapid correlated evolutionary adaptation. Behavioral Ecology. 23(5):960–969. doi:10.1093/beheco/ars059.

Atwell JW, Cardoso GC, Whittaker DJ, Price TD, Ketterson ED. 2014. Hormonal, behavioral, and life-history traits exhibit correlated shifts in relation to population establishment in a novel environment. The American Naturalist. 184(6):E147–E160. doi:10.1086/678398.

Baxter-Gilbert J, Riley JL, Whiting MJ. 2019. Bold new world: urbanization promotes an innate behavioral trait in a lizard. Behavioral Ecology and Sociobiology. 73(8):1–10.

Biard C, Brischoux F, Meillère A, Michaud B, Nivière M, Ruault S, Vaugoyeau M, Angelier F. 2017. Growing in cities: An urban penalty for wild birds? A study of phenotypic differences between urban and rural great tit chicks (*Parus major*). Frontiers in Ecology and Evolution. 5:1–14. doi:10.3389/fevo.2017.00079.

Bókony V, Balogh E, Ujszegi J, Ujhegyi N, Szederkényi M, Hettyey A. 2024. Tadpoles develop elevated heat tolerance in urban heat islands regardless of sex. Evol Biol. 51(1):209–216. doi:10.1007/s11692-024-09626-7.

Bókony V, Ujhegyi N, Hamow KÁ, Bosch J, Thumsová B, Vörös J, Aspbury AS, Gabor CR. 2021. Stressed tadpoles mount more efficient glucocorticoid negative feedback in anthropogenic habitats due to phenotypic plasticity. Science of the Total Environment. 753:141896. doi:10.1016/j.scitotenv.2020.141896.

Bókony V, Üveges B, Verebélyi V, Ujhegyi N, Móricz ÁM. 2019. Toads phenotypically adjust their chemical defences to anthropogenic habitat change. Scientific reports. 9(1):1–8.

Bonier F. 2012. Hormones in the city: Endocrine ecology of urban birds. Hormones and Behavior. 61(5):763–772. doi:10.1016/j.yhbeh.2012.03.016.

Brans KI, Almeida RA, Fajgenblat M. 2021. Genetic differentiation in pesticide resistance between urban and rural populations of a nontarget freshwater keystone interactor, *Daphnia magna*. Evolutionary Applications. 14(10):2541–2552. doi:10.1111/eva.13293.

Brans KI, De Meester L. 2018. City life on fast lanes: Urbanization induces an evolutionary shift towards a faster lifestyle in the water flea *Daphnia*. Functional Ecology. 32(9):2225–2240.

Brans KI, Govaert L, Engelen JMT, Gianuca AT, Souffreau C, De Meester L. 2017. Eco-evolutionary dynamics in urbanized landscapes: evolution, species sorting and the change in zooplankton body size along urbanization gradients. Philosophical Transactions of the Royal Society B: Biological Sciences. 372(1712):20160030.

Brans KI, Jansen M, Vanoverbeke J, Tüzün N, Stoks R, De Meester L. 2017. The heat is on: Genetic adaptation to urbanization mediated by thermal tolerance and body size. Global change biology. 23(12):5218–5227.

Brans KI, Stoks R, De Meester L. 2018. Urbanization drives genetic differentiation in physiology and structures the evolution of pace-of-life syndromes in the water flea *Daphnia magna*. Proceedings of the Royal Society B: Biological Sciences. 285(1883). doi:10.1098/rspb.2018.0169.

Brans KI, Tüzün N, Sentis A, De Meester L, Stoks R. 2022. Cryptic eco evolutionary feedback in the city: Urban evolution of prey dampens the effect of urban evolution of the predator. Journal of Animal Ecology. 91(3):514–526. doi:10.1111/1365-2656.13601.

Breitbart ST, Agrawal AA, Wagner HH, Johnson MT. 2023. Urbanization and a green corridor do not impact genetic divergence in common milkweed (*Asclepias syriaca L*.). Scientific Reports. 13(1):20437.

Caizergues AE, Charmantier A, Lambrechts MM, Perret S, Demeyrier V, Lucas A, Grégoire A. 2021. An avian urban morphotype: how the city environment shapes great tit morphology at different life stages. Urban Ecosystems. 45(5):929–941. doi:10.1007/s11252-020-01077-0.

Caizergues AE, Grégoire A, Charmantier A. 2018. Urban versus forest ecotypes are not explained by divergent reproductive selection. Proceedings of the Royal Society B: Biological Sciences. 285:20180261.

Caizergues AE, Grégoire A, Choquet R, Perret S, Charmantier A. 2022. Are behaviour and stress-related phenotypes in urban birds adaptive? Journal of Animal Ecology. 91(8):1627–1641.

Caizergues AE, Le Luyer J, Grégoire A, Szulkin M, Senar J, Charmantier A, Perrier C. 2022. Epigenetics and the city: Non parallel DNA methylation modifications across pairs of urban forest Great tit populations. Evolutionary Applications. 15(1):149–165. doi:10.1111/eva.13334.

Caizergues AE, Robira B, Perrier C, Jeanneau M, Berthomieu A, Perret S, Gandon S, Charmantier A. 2024. Cities as parasitic amplifiers? Malaria prevalence and diversity in great tits along an urbanization gradient. Peer Community Journal. 4:article no. e38. doi:10.24072/pcjournal.405.

Camacho C, Sanabria-Fernández A, Baños-Villalba A, Edelaar P. 2020. Experimental evidence that matching habitat choice drives local adaptation in a wild population. Proceedings of the Royal Society B: Biological Sciences. 287:20200721.

Campbell-Staton SC, Velotta JP, Winchell KM. 2021. Selection on adaptive and maladaptive gene expression plasticity during thermal adaptation to urban heat islands. Nature communications. 12(1):6195.

Capilla-Lasheras P, Thompson MJ, Sánchez-Tójar A, Haddou Y, Branston CJ, Réale D, Charmantier A, Dominoni DM. 2022. A global meta-analysis reveals higher variation in breeding phenology in urban birds than in their non-urban neighbours. Ecology Letters. 25(11):2552–2570.

Carere C, van Oers K. 2004. Shy and bold great tits (*Parus major*): Body temperature and breath rate in response to handling stress. Physiology and Behavior. 82(5):905–912. doi:10.1016/j.physbeh.2004.07.009.

Caspi T, Johnson JR, Lambert MR, Schell CJ, Sih A. 2022. Behavioral plasticity can facilitate evolution in urban environments. Trends in Ecology & Evolution. 37(12):1092–1103.

Charmantier A, Demeyrier V, Lambrechts M, Perret S, Grégoire A. 2017. Urbanization is associated with divergence in pace-of-life in great tits. Frontiers in Ecology and Evolution. 5:1–13. doi:10.3389/fevo.2017.00053.

Charmantier A, Garant D, Kruuk LE. 2014. Quantitative genetics in the wild. OUP Oxford.

Cheptou P-O, Carrue O, Rouifed S, Cantarel A. 2008. Rapid evolution of seed dispersal in an urban environment in the weed Crepis sancta. Proceedings of the National Academy of Sciences. 105(10):3796–3799.

Chick LD, Strickler SA, Perez A, Martin RA, Diamond SE. 2019. Urban heat islands advance the timing of reproduction in a social insect. Journal of Thermal Biology. 80:119–125. doi:10.1016/j.jtherbio.2019.01.004.

Chick LD, Waters JS, Diamond SE. 2021. Pedal to the metal: Cities power evolutionary divergence by accelerating metabolic rate and locomotor performance. Evolutionary Applications. 14(1):36–52. doi:10.1111/eva.13083.

Christians JK. 2002. Avian egg size: variation within species and inflexibility within individuals. Biological reviews. 77(1):1–26.

Conover DO, Duffy TA, Hice LA. 2009. The covariance between genetic and environmental influences across ecological gradients. The Year in Evolutionary Biology 2009, Volume 1168. 28:100.

Corsini M, Schöll EM, Di Lecce I, Chatelain M, Dubiec A, Szulkin M. 2021. Growing in the city: Urban evolutionary ecology of avian growth rates. Evolutionary Applications. 14(1):69–84. doi:10.1111/eva.13081.

Czaczkes TJ, Bastidas-Urrutia AM, Ghislandi P, Tuni C. 2018. Reduced light avoidance in spiders from populations in light-polluted urban environments. The Science of Nature. 105(11):1–5.

Demeyrier V, Lambrechts MM, Perret P, Grégoire A. 2016. Experimental demonstration of an ecological trap for a wild bird in a human-transformed environment. Animal Behaviour. 118:181–190. doi:10.1016/j.anbehav.2016.06.007.

Des Roches S, Pendleton LH, Shapiro B, Palkovacs EP. 2021. Conserving intraspecific variation for nature’s contributions to people. Nature Ecology and Evolution. 5(5):574–582. doi:10.1038/s41559-021-01403-5.

Dhondt AA. 1982. Heritability of blue tit tarsus length from normal and cross-fostered broods. Evolution. 36(2):418–419.

Diamond SE, Chick L, Perez ABE, Strickler SA, Martin RA. 2017. Rapid evolution of ant thermal tolerance across an urban-rural temperature cline. Biological Journal of the Linnean Society. 121(2):248–257.

Diamond Sarah E., Chick LA, Perez A, Strickler SA, Zhao C. 2018. Evolution of plasticity in the city: Urban acorn ants can better tolerate more rapid increases in environmental temperature. Conservation Physiology. 6(1):1–12. doi:10.1093/conphys/coy030.

Diamond Sarah E, Chick LD, Perez A, Strickler SA, Martin RA. 2018. Evolution of thermal tolerance and its fitness consequences: parallel and non-parallel responses to urban heat islands across three cities. Proceedings of the Royal Society B: Biological Sciences. 285(1882):20180036.

Diamond SE, Martin RA. 2021. Evolution in cities. Annual Review of Ecology, Evolution, and Systematics. 52:519–540.

Diamond SE, Martin RA, Bellino G, Crown KN, Prileson EG. 2022. Urban evolution of thermal physiology in a range-expanding, mycophagous fruit fly, *Drosophila tripunctata*. Biological Journal of the Linnean Society. 137(3):409–420.

Dubuc-Messier G, Caro SP, Perrier C, van Oers K, Réale D, Charmantier A. 2018. Gene flow does not prevent personality and morphological differentiation between two blue tit populations. Journal of Evolutionary Biology. 31(8):1127–1137. doi:10.1111/jeb.13291.

Dubuc-Messier G, Réale D, Perret P, Charmantier A. 2016. Environmental heterogeneity and population differences in blue tits personality traits. Behavioral Ecology. 28:arw148. doi:10.1093/beheco/arw148.

Edelaar P, Siepielski AM, Clobert J. 2008. Matching habitat choice causes directed gene flow: A neglected dimension in evolution and ecology. Evolution. 62(10):2462–2472. doi:10.1111/j.1558-5646.2008.00459.x.

Fudickar AM, Greives TJ, Abolins-Abols M, Atwell JW, Meddle SL, Friis G, Stricker CA, Ketterson ED. 2017. Mechanisms associated with an advance in the timing of seasonal reproduction in an urban songbird. Frontiers in Ecology and Evolution. 5:85.

Fukano Y, Guo W, Uchida K, Tachiki Y. 2020. Contemporary adaptive divergence of plant competitive traits in urban and rural populations and its implication for weed management. Journal of Ecology. 108(6):2521–2530.

Gebhardt-Henrich SG, Van Noordwijk AJ. 1991. Nestling growth in the great tit I. Heritability estimates under different environmental conditions. Journal of Evolutionary Biology. 4(3):341–362.

Géron C, Lembrechts JJ, Hamdi R, Berckmans J, Nijs I, Monty A. 2022. Phenotypic variation along urban-to-rural gradients: an attempt to disentangle the mechanisms at play using the alien species Matricaria discoidea (Asteraceae). Plant Ecology. 223(10):1219–1231.

Ghalambor CK, McKay JK, Carroll SP, Reznick DN. 2007. Adaptive versus non-adaptive phenotypic plasticity and the potential for contemporary adaptation in new environments. Functional Ecology. 21(3):394–407. doi:10.1111/j.1365-2435.2007.01283.x.

y Gomez GSM, Van Dyck H. 2012. Ecotypic differentiation between urban and rural populations of the grasshopper Chorthippus brunneus relative to climate and habitat fragmentation. Oecologia. 169(1):125–133.

Gorton AJ, Moeller DA, Tiffin P. 2018. Little plant, big city: a test of adaptation to urban environments in common ragweed (*Ambrosia artemisiifolia*). Proceedings of the Royal Society B: Biological Sciences. 285(1881):20180968.

Gosler AG, Harper DGC. 2000. Assessing the heritability of body condition in birds: a challenge exemplified by the great tit Parus major L.(Aves). Biological Journal of the Linnean Society. 71(1):103–117.

Hadfield JD. 2010. MCMC Methods for Multi-Response Generalized Linear Mixed Models: The {MCMCglmm} {R} Package. Journal of Statistical Software. 33(2):1–22.

Hahs AK, Fournier B, Aronson MF, Nilon CH, Herrera-Montes A, Salisbury AB, Threlfall CG, Rega-Brodsky CC, Lepczyk CA, La Sorte FA. 2023. Urbanisation generates multiple trait syndromes for terrestrial animal taxa worldwide. Nature communications. 14(1):4751.

Harten L, Gonceer N, Handel M, Dash O, Fokidis HB, Yovel Y. 2021. Urban bat pups take after their mothers and are bolder and faster learners than rural pups. BMC Biol. 19(1):190. doi:10.1186/s12915-021-01131-z.

Hendry AP. 2017. Eco-evolutionary Dynamics. Princeton: Princeton University Press. [accessed 2024 Apr 12]. https://www.degruyter.com/document/doi/10.1515/9781400883080/html.

Hendry AP, Farrugia TJ, Kinnison MT. 2008. Human influences on rates of phenotypic change in wild animal populations. Molecular Ecology. 17(1):20–29. doi:10.1111/j.1365-294X.2007.03428.x.

Hõrak P. 1994. Effect of nestling history on adult size and reproduction in the Great Tit. Ornis fennica. 71(2):47–54.

Hõrak P, Mänd R, Ots I, Leivits A. 1995. Egg size in the Great Tit Parus major: individual, habitat and geographic differences. Ornis fennica. 72(3):97–114.

Hwang CC, Turner BD. 2009. Small scaled geographical variation in life history traits of the blowfly *Calliphora vicina* between rural and urban populations. Entomologia Experimentalis et Applicata. 132(3):218–224.

Ichikawa I, Kuriwada T. 2023. The combined effects of artificial light at night and anthropogenic noise on life history traits in ground crickets. Ecological Research. 38(3):446–454. doi:10.1111/1440-1703.12380.

Irwin RE, Warren PS, Carper AL, Adler LS. 2014. Plant–animal interactions in suburban environments: implications for floral evolution. Oecologia. 174:803–815.

Jacquier L., Doums C, Four-Chaboussant A, Peronnet R, Tirard C, Molet M. 2021. Urban colonies are more resistant to a trace metal than their forest counterparts in the ant *Temnothorax nylanderi*. Urban Ecosystems. 24:561–570.

Jacquier Lauren, Molet M, Bocquet C, Doums C. 2021. Hibernation conditions contribute to the differential resistance to cadmium between urban and forest ant colonies. Animals. 11(4):1050.

Kaiser A, Merckx T, Van Dyck H. 2018. Urbanisation and sex affect the consistency of butterfly personality across metamorphosis. Behavioral Ecology and Sociobiology. 72(12):1–11.

Kaiser A, Merckx T, Van Dyck H. 2020. An experimental test of changed personality in butterflies from anthropogenic landscapes. Behavioral Ecology and Sociobiology. 74(7):1–8.

Kawecki TJ, Ebert D. 2004. Conceptual issues in local adaptation. Ecology letters. 7(12):1225–1241.

Kern E, Langerhans RB. 2019. Urbanization alters swimming performance of a stream fish. Frontiers in Ecology and Evolution. 6:229.

Kern EMA, Langerhans RB. 2018. Urbanization drives contemporary evolution in stream fish. Global Change Biology. 24(8):3791–3803.

Kinnison MT, Hendry AP. 2001. The pace of modern life II: From rates of contemporary microevolution to pattern and process. Genetica. 112–113:145–164. doi:10.1007/978-94-010-0585-2_10.

Koolhaas JM, Bartolomucci A, Buwalda B, de Boer SF, Flügge G, Korte SM, Meerlo P, Murison R, Olivier B, Palanza P. 2011. Stress revisited: a critical evaluation of the stress concept. Neuroscience & Biobehavioral Reviews. 35(5):1291–1301.

Koolhaas JM, De Boer SF, Coppens CM, Buwalda B. 2010. Neuroendocrinology of coping styles: towards understanding the biology of individual variation. Frontiers in neuroendocrinology. 31(3):307–321.

Kostanecki A, Gorton AJ, Moeller DA. 2021. An urban–rural spotlight: Evolution at small spatial scales among urban and rural populations of common ragweed. Journal of Urban Ecology. 7(1):juab004.

Krams IA, Vrublevska J, Sepp T, Abolins Abols M, Rantala MJ, Mierauskas P, Krama T. 2014. Sex specific associations between nest defence, exploration and breathing rate in breeding pied flycatchers. Wright J, editor. Ethology. 120(5):492–501. doi:10.1111/eth.12222.

Kuriwada T. 2023. Differences in male calling song and female mate location behaviour between urban and rural crickets. Biological Journal of the Linnean Society. 139(3):275–285.

Lambert MR, Brans KI, Roches SD, Donihue CM, Diamond SE. 2021. Adaptive evolution in cities: progress and misconceptions. Trends in Ecology & Evolution. 36(3):239–257.

Lambert MR, Donihue CM. 2020. Urban biodiversity management using evolutionary tools. Nature Ecology and Evolution. 4(7):903–910. doi:10.1038/s41559-020-1193-7.

Lambrecht SC, Mahieu S, Cheptou P-O. 2016. Natural selection on plant physiological traits in an urban environment. Acta Oecologica. 77:67–74.

Lampe U, Reinhold K, Schmoll T. 2014. How grasshoppers respond to road noise: Developmental plasticity and population differentiation in acoustic signalling. Functional Ecology. 28(3):660–668. doi:10.1111/1365-2435.12215.

Leinonen T, Mccairns RJS, Hara RBO, Merilä J. 2013. QST – FST comparisons : evolutionary and ecological insights from genomic heterogeneity. Nature Reviews Genetics. doi:10.1038/nrg3395.

Martin RA, Chick LD, Yilmaz AR, Diamond SE. 2019. Evolution, not transgenerational plasticity, explains the adaptive divergence of acorn ant thermal tolerance across an urban–rural temperature cline. Evolutionary applications. 12(8):1678–1687.

McLean MA, Angilletta Jr MJ, Williams KS. 2005. If you can’t stand the heat, stay out of the city: thermal reaction norms of chitinolytic fungi in an urban heat island. Journal of Thermal Biology. 30(5):384–391.

Merckx T, Nielsen ME, Heliölä J, Kuussaari M, Pettersson LB, Pöyry J, Tiainen J, Gotthard K, Kivelä SM. 2021. Urbanization extends flight phenology and leads to local adaptation of seasonal plasticity in Lepidoptera. Proc Natl Acad Sci USA. 118(40):e2106006118. doi:10.1073/pnas.2106006118.

Merckx T, Nielsen ME, Kankaanpää T, Kadlec T, Yazdanian M, Kivelä SM. 2023. Dim light pollution prevents diapause induction in urban and rural moths. Journal of Applied Ecology. 60(6):1022–1031. doi:10.1111/1365-2664.14373.

Merckx T, Nielsen ME, Kankaanpää T, Kadlec T, Yazdanian M, Kivelä SM. 2024. Continent wide parallel urban evolution of increased heat tolerance in a common moth. Evolutionary Applications. 17(1):e13636. doi:10.1111/eva.13636.

Merckx T, Souffreau C, Kaiser A, Baardsen LF, Backeljau T, Bonte D, Brans KI, Cours M, Dahirel M, Debortoli N, et al. 2018. Body-size shifts in aquatic and terrestrial urban communities. Nature. 558(7708):113–116. doi:10.1038/s41586-018-0140-0.

Merilä J, Fry JD. 1998. Genetic variation and causes of genotype-environment interaction in the body size of blue tit (*Parus caeruleus*). Genetics. 148(3):1233–1244.

Mertens JAL. 1977. Thermal conditions for successful breeding in great tits (Parus major L.) I. Relation of growth and development of temperature regulation in nestling great tits. Oecologia. 28(1):1–29.

Miranda AC, Schielzeth H, Sonntag T, Partecke J. 2013. Urbanization and its effects on personality traits: A result of microevolution or phenotypic plasticity? Global Change Biology. 19(9):2634–2644. doi:10.1111/gcb.12258.

Mitchell-Olds T, Willis JH, Goldstein DB. 2007. Which evolutionary processes influence natural genetic variation for phenotypic traits? Nature Reviews. 8:845–856. doi:10.1038/nrg2207.

Mohajerani A, Bakaric J, Jeffrey-Bailey T. 2017. The urban heat island effect, its causes, and mitigation, with reference to the thermal properties of asphalt concrete. Journal of environmental management. 197:522–538.

Mühlenhaupt M, Baxter-Gilbert J, Makhubo BG, Riley JL, Measey J. 2022. No evidence for innate differences in tadpole behavior between natural, urbanized, and invasive populations. Behavioral Ecology Sociobiology. 76(1):11. doi:10.1007/s00265-021-03121-1.

Nicolaus M, Edelaar P. 2018. Comparing the consequences of natural selection, adaptive phenotypic plasticity, and matching habitat choice for phenotype–environment matching, population genetic structure, and reproductive isolation in meta-populations. Ecology and Evolution. 8(8):3815–3827. doi:10.1002/ece3.3816.

Palomar G, Wos G, Stoks R, Sniegula S. 2023. Latitude specific urbanization effects on life history traits in the damselfly *Ischnura elegans*. Evolutionary Applications. 16(8):1503–1515. doi:10.1111/eva.13583.

Partecke J, Gwinner E. 2007. Increased sedentariness in European blackbirds following urbanization: a consequence of local adaptation? Ecology. 88(4):882–890.

Partecke J, Schwabl I, Gwinner E. 2006. Stress and the city: Urbanization and its effects on the stress physiology in European blackbirds. Ecology. 87(8):1945–1952. doi:10.1890/0012-9658(2006)87[1945:SATCUA]2.0.CO;2.

Perrier C, Lozano del Campo A, Szulkin M, Demeyrier V, Gregoire A, Charmantier A. 2018. Great tits and the city: Distribution of genomic diversity and gene–environment associations along an urbanization gradient. Evolutionary Applications. 11(5):593–613. doi:10.1111/eva.12580.

Pipoly I, Preiszner B, Sándor K, Sinkovics C, Seress G, Vincze E, Bókony V, Liker A. 2022. Extreme hot weather has stronger impacts on Avian reproduction in forests than in cities. Frontiers in Ecology and Evolution. 10.

Pisman M, Bonte D, De La Peña E. 2020. Urbanization alters plastic responses in the common dandelion *Taraxacum officinale*. Ecology and Evolution. 10(9):4082–4090. doi:10.1002/ece3.6176.

Playà Montmany N, González Medina E, Cabello Vergel J, Parejo M, Abad Gómez JM, Sánchez Guzmán JM, Villegas A, Masero JA. 2021. The thermoregulatory role of relative bill and leg surface areas in a Mediterranean population of Great tit (*Parus major*). Ecology and Evolution. 11(22):15936–15946. doi:10.1002/ece3.8263.

Qu J, Bonte D, Vandegehuchte ML. 2022. Phenotypic and genotypic divergence of plant– herbivore interactions along an urbanization gradient. Evolutionary Applications. 15(5):865–877. doi:10.1111/eva.13376.

R Core Team. 2024. R: A language and environment for statistical computing.

Rasner CA, Yeh P, Eggert LS, Hunt KE, Woodruff DS, Price TD. 2004. Genetic and morphological evolution following a founder event in the dark eyed junco, *Junco hyemalis thurberi*. Molecular Ecology. 13(3):671–681.

Reichard DG, Atwell JW, Pandit MM, Cardoso GC, Price TD, Ketterson ED. 2020. Urban birdsongs: higher minimum song frequency of an urban colonist persists in a common garden experiment. Animal Behaviour. 170:33–41. doi:10.1016/j.anbehav.2020.10.007.

Rivkin LR, Santangelo JS, Alberti M, Aronson MFJ, de Keyzer CW, Diamond SE, Fortin MJ, Frazee LJ, Gorton AJ, Hendry AP, et al. 2019. A roadmap for urban evolutionary ecology. Evolutionary Applications. 12(3):384–398. doi:10.1111/eva.12734.

Salmón P, Jacobs A, Ahrén D, Biard C, Dingemanse NJ, Dominoni DM, Helm B, Lundberg M, Senar JC, Sprau P, et al. 2021. Continent-wide genomic signatures of adaptation to urbanisation in a songbird across Europe. Nature Communications. 12(1):2983. doi:10.1038/s41467-021-23027-w.

Samocha Y, Scharf I. 2020. Comparison of wormlion behavior under man-made and natural shelters: urban wormlions more strongly prefer shaded, fine-sand microhabitats, construct larger pits and respond faster to prey. Current Zoology. 66(1):91–98.

Sanderson S, Bolnick DI, Kinnison MT, O’Dea RE, Gorné LD, Hendry AP, Gotanda KM. 2023. Contemporary changes in phenotypic variation, and the potential consequences for eco evolutionary dynamics. Ecology Letters. 26(S1). doi:10.1111/ele.14186.

Santangelo JS, Miles LS, Breitbart ST, Murray-Stoker D, Rivkin LR, Johnson MTJ, Ness RW. 2020. Urban environments as a framework to study parallel evolution. In: Szulkin M, Munshi-South J, Charmantier A, editors. Urban Evolutionary Biology. USA: Oxford University Press. p. 36–53.

Sato A, Takahashi Y. 2022. Responses in thermal tolerance and daily activity rhythm to urban stress in *Drosophila suzukii*. Ecology and Evolution. 12(12):e9616. doi:10.1002/ece3.9616.

Schmitz G, Linstädter A, Frank ASK, Dittberner H, Thome J, Schrader A, Linne von Berg K, Fulgione A, Coupland G, De Meaux J. 2024. Environmental filtering of life history trait diversity in urban populations of *Arabidopsis thaliana*. Journal of Ecology. 112(1):14–27. doi:10.1111/1365-2745.14211.

Seress G, Sándor K, Evans KL, Liker A. 2020. Food availability limits avian reproduction in the city: An experimental study on great tits *Parus major*. Journal of Animal Ecology. 89(7):1570–1580. doi:10.1111/1365-2656.13211.

Snell-Rood EC. 2013. An overview of the evolutionary causes and consequences of behavioural plasticity. Animal Behaviour. 85(5):1004–1011. doi:10.1016/j.anbehav.2012.12.031.

Sol D, Lapiedra O, González-Lagos C. 2013. Behavioural adjustments for a life in the city. Animal Behaviour. 85(5):1101–1112. doi:10.1016/j.anbehav.2013.01.023.

Stirling DG, Réale D, Roff DA. 2002. Selection, structure and the heritability of behaviour. Journal of Evolutionary Biology. 15(2):277–289.

Sumasgutner P, Cunningham SJ, Hegemann A, Amar A, Watson H, Nilsson JF, Andersson MN, Isaksson C. 2023. Interactive effects of rising temperatures and urbanisation on birds across different climate zones: A mechanistic perspective. Global Change Biology. 29(9):2399–2420. doi:10.1111/gcb.16645.

Svensson L. 1992. Identification guide to European passerines. The author.

Szulkin M, Munshi-South J, Charmantier A. 2020. Urban evolutionary biology. Oxford University Press, USA.

Taichi N, Ushimaru A. 2024. Trait variation along an urban–rural gradient in Asian dayflower: the contribution of phenotypic plasticity and genetic divergence. Plant Biol J. 26(1):74–81. doi:10.1111/plb.13595.

Talloen W, Lens LUC, Van Dongen S, Adriaensen F, Matthysen E. 2010. Mild stress during development affects the phenotype of great tit *Parus major* nestlings: a challenge experiment. Biological Journal of the Linnean society. 100(1):103–110.

Tene Fossog B, Antonio-Nkondjio C, Kengne P, Njiokou F, Besansky NJ, Costantini C. 2013. Physiological correlates of ecological divergence along an urbanization gradient: differential tolerance to ammonia among molecular forms of the malaria mosquito *Anopheles gambiae*. BMC ecology. 13(1):1–12.

Tene Fossog B, Poupardin R, Costantini C, Awono-Ambene P, Wondji CS, Ranson H, Antonio-Nkondjio C. 2013. Resistance to DDT in an urban setting: common mechanisms implicated in both M and S forms of *Anopheles gambiae* in the city of Yaoundé Cameroon. PloS one. 8(4):e61408.

Thompson KA, Renaudin M, Johnson MTJ. 2016. Urbanization drives the evolution of parallel clines in plant populations. Proceedings of the Royal Society B: Biological Sciences. 283(1845). doi:10.1098/rspb.2016.2180.

Thompson MJ, Capilla-Lasheras P, Dominoni DM, Réale D, Charmantier A. 2022. Phenotypic variation in urban environments: mechanisms and implications. Trends in Ecology & Evolution. 37:171–182.

Tüzün N, Debecker S, de Beeck LO, Stoks R. 2015. Urbanisation shapes behavioural responses to a pesticide. Aquatic Toxicology. 163:81–88.

Tüzün N, Mueller S, Koch K, Stoks R. 2017. Pesticide-induced changes in personality depend on the urbanization level. Animal behaviour. 134:45–55.

Tüzün N, Op de Beeck L, Brans KI, Janssens L, Stoks R. 2017. Microgeographic differentiation in thermal performance curves between rural and urban populations of an aquatic insect. Evolutionary applications. 10(10):1067–1075.

Tüzün N, Stoks R. 2021. Lower bioenergetic costs but similar immune responsiveness under a heat wave in urban compared to rural damselflies. Evolutionary applications. 14(1):24–35.

Van de Schoot E, Merckx T, Ebert D, Wesselingh RA, Altermatt F, Van Dyck H. 2024. Evolutionary change in flight-to-light response in urban moths comes with changes in wing morphology. Biol Lett. 20(3):20230486. doi:10.1098/rsbl.2023.0486.

Van Noordwijk AJ, Keizer LCP, Van Balen JH, Scharloo W. 1981. Genetic variation in egg dimensions in natural populations of the Great Tit. Genetica. 55:221–232.

Vaugoyeau M, Adriaensen F, Artemyev A, Bańbura J, Barba E, Biard C, Blondel J, Bouslama Z, Bouvier J-C, Camprodon J, et al. 2016. Interspecific variation in the relationship between clutch size, laying date and intensity of urbanization in four species of hole-nesting birds. Ecology and Evolution. 6(16):5907–5920. doi:10.1002/ece3.2335.

de Villemereuil P. 2018. Quantitative genetic methods depending on the nature of the phenotypic trait. Annals of the New York Academy of Sciences. 1422(1):29–47. doi:10.1111/nyas.13571.

de Villemereuil P, Gaggiotti OE, Goudet J. 2020. Common garden experiments to study local adaptation need to account for population structure. Journal of Ecology. 00.

de Villemereuil P, Morrissey MB, Nakagawa S, Schielzeth H. 2018. Fixed-effect variance and the estimation of repeatabilities and heritabilities: Issues and solutions. Journal of Evolutionary Biology. 31(4):621–632.

de Villemereuil P, Mouterde M, Gaggiotti OE, Till Bottraud I. 2018. Patterns of phenotypic plasticity and local adaptation in the wide elevation range of the alpine plant *Arabis alpina*. Jacquemyn H, editor. Journal of Ecology. 106(5):1952–1971. doi:10.1111/1365-2745.12955.

de Villemereuil P, Schielzeth H, Nakagawa S, Morrissey M. 2016. General methods for evolutionary quantitative genetic inference from generalized mixed models. Genetics. 204(3):1281–1294. doi:10.1534/genetics.115.186536.

Westby KM, Medley KA. 2020. Cold nights, city lights: artificial light at night reduces photoperiodically induced diapause in urban and rural populations of *aedes albopictus* (Diptera: *Culicidae*). Journal of Medical Entomology. 57(6):1694–1699.

Weston LM, Mattingly KZ, Day CTC, Hovick SM. 2021. Potential local adaptation in populations of invasive reed canary grass (*Phalaris arundinacea*) across an urbanization gradient. Ecology and Evolution. 11(16):11457–11476. doi:10.1002/ece3.7938.

Whitehead A, Pilcher W, Champlin D, Nacci D. 2012. Common mechanism underlies repeated evolution of extreme pollution tolerance. Proceedings of the Royal Society B: Biological Sciences. 279(1728):427–433.

Wilson AJ. 2018. How should we interpret estimates of individual repeatability? Evolution Letters.:1–5. doi:10.1002/evl3.40.

Winchell KM, Reynolds RG, Prado Irwin SR, Puente Rolón AR, Revell LJ. 2016. Phenotypic shifts in urban areas in the tropical lizard Anolis cristatellus. Evolution. 70(5):1009–1022.

Yakub M, Tiffin P. 2017. Living in the city: urban environments shape the evolution of a native annual plant. Global change biology. 23(5):2082–2089.

Yeh PJ. 2004. Rapid evolution of a sexually selected trait following population establishment in a novel habitat. Evolution. 58(1):166–174.

Yeh PJ, Price TD. 2004. Adaptive Phenotypic Plasticity and the Successful Colonization of a Novel Environment. American Naturalist. 164(4):531–542.

Yilmaz AR, Diamond SE, Martin RA. 2021. Evidence for the evolution of thermal tolerance, but not desiccation tolerance, in response to hotter, drier city conditions in a cosmopolitan, terrestrial isopod. Evolutionary applications. 14(1):12–23.

Young EA, Postma E. 2023. Low interspecific variation and no phylogenetic signal in additive genetic variance in wild bird and mammal populations. Ecology and Evolution. 13(11):e10693. doi:10.1002/ece3.10693.

Youngflesh C, Saracco JF, Siegel RB, Tingley MW. 2022. Abiotic conditions shape spatial and temporal morphological variation in North American birds. Nature Ecology & Evolution. 6(12):1860–1870.

Zhang H, He Y, Yang J, Mao H, Jiang X. 2022. Contemporary adaptive evolution in fragmenting river landscapes: evidence from the native waterflea Ceriodaphnia cornuta. Journal of Plankton Research. 44(1):88–98.

