## Supplementary materials for "The city and forest bird flock together in a common garden: genetic and environmental effects drive urban phenotypic divergence"

*Caro and Charmantier should be considered joint last authors

[1] Département des sciences biologiques, Université du Québec à Montréal, 141 Avenue du Président-Kennedy, Montréal, QC H2X 1Y4, Canada

[2] CEFE, Univ Montpellier, CNRS, EPHE, IRD, Montpellier, FRANCE

[3] Parc de Lunaret, Ville de Montpellier, 50 Avenue Agropolis, 34090, Montpellier, France

141 Av. du Président-Kennedy, Montréal, QC, H2X 1Y4, Canada

Supplementary methods:

*Quantifying urbanisation*

We quantified urbanization at each nest box using impervious surface area (ISA; sealed non-natural surfaces) from the Copernicus imperviousness density raster data set (resolution 10 m, tiles: E38N22/E38N23, projection: LAEA EPSG 3035; European Environment Agency 2020). We calculated the proportion of ISA using a 100-meter radius circular buffer around each nest box in QGIS (v3.22.0; QGIS Development Team 2023) and calculated a ratio between the number of ISA pixels over the number of pixels contained in the buffer area (range = 0 – 1, where 1 is all ISA). We used the proportion ISA at each nest box as a continuous urbanization metric to characterize the territory of breeding wild birds (captured at nest boxes) and the territory of origin for the birds raised during the common garden experiment.

*Captive diet*

During the nestling stage, diet consisted of hand-rearing powder solution (Nutribird A21 and A19, Versele-Laga, Deinze, Belgium), alternating with dead wax moth larvae and mealworms. Once chicks were 15 days old, the diet was enriched with a cake made of eggs, sunflower margarine, sugar, wheat and protein-rich pellet flours (Country's Best Show1-2 Crumble, Versele-Laga, Deinze, Belgium). Cake was supplemented with commercial powders containing mostly vitamins and minerals (Nutribird A21, Versele-Laga; and Nekton-S, Nekton GmbH, Pforzheim, Germany). After transfer to outdoor aviaries, birds were eating independently a diet made of cake (see above) and live mealworms. Food and water were provided *ad libitum*.

*Genotyping*

We extracted DNA from blood samples using DNeasy blood and tissue kits. Extracted DNA was sent to the Montpellier GenomiX platform (MGX) for RAD sequencing following a similar protocol as described in (Caizergues et al. 2022). We had a total of N = 343 individual samples for sequencing (N = 270 wild and 73 common garden individuals), which generated 10.1 M reads and an average depth of 19.8x per individual using paired-end RAD sequencing (2*150pb). We obtained 185321 SNPs on autosomal chromosome after filtering and we randomly subsampled 600 independent SNPs with a minor Allele Frequency (MAF) of 0.4 to reconstruct the genetic relatedness between all birds (using R package 'Sequoia'; Huisman 2017).

In one foster brood, we could identify relatedness between individuals, but not their origin nest as we were missing information on parental identity (Table S2; foster nest ZOO46 received nestlings from FAC2 and CEF7). As we could not identify the urbanization level of the origin habitat with certainty for these cases, we used the average proportion ISA between the possible origin nest boxes as their urbanization value. Genotyping revealed that the ZOO46 foster nest contained three pairs of siblings (i.e., 6 individuals with 3 unique mothers and fathers). This was presumably because the four eggs collected from origin nest CEF7 contained two separate pairs of siblings; most likely a case of egg dumping. Therefore, we coded individuals from foster nest ZOO46 as being from three different origin nests to account for relatedness between individuals and differences in maternal egg investment.

*Full description of wild and common garden mixed models*

*Aggression in hand:* We fit aggression in hand scores as the response variable using Gaussian mixed-effect models. On average, we measured individuals 1.69 times in the wild (range: 1 – 8) and 3.89 times in the common garden (range: 1 – 4; approx. 44 to 264 days old; Table S2). For the wild model, we also controlled for age category (adult vs. yearling), the time of day in continuous format (minutes divided by 60), and the Julian date of measure (days since Jan 1) as fixed effects. For the CG model, we fit the fixed effects of time of day in continuous format and the assay number. Assay number in the CG combines age effects, date or seasonal effects, and habituation effects simultaneously as these factors are correlated with repeated assays over time in our experiment. In the case of aggression in hand, assay number also accounted for observer effects as observer ID was confounded with assay number and could not be estimated for this trait (i.e., only a single observer scored first aggression assay). Results were qualitatively similar when accounting for observer ID as a fixed effect in these models instead of assay number.

*Breath rate index:* We fit breath rate index as the response variable in Gaussian mixed-effect models. On average, we measured individuals 1.32 times in the wild (range: 1 – 6) and 3.89 times in the common garden (range: 1 – 4; approx. 44 to 264 days old; Table S2). For the wild model, we also fit age category, Julian date of measure, and protocol type (new vs. old) as fixed effects. Initially breath rate index was measured as the number of breaths for 30 seconds (between 2013 – 2016) and so we account for this difference in protocol since these initial measures have been converted to approximate the amount of time for 30 breaths (Caizergues et al. 2022). As time of day and temperature were correlated in this data set ($r^2 = 0.48$, $P < 0.001$), we chose to only include temperature as an additional fixed effect since this variable explained more variation in this trait than time of day. For the CG model, we fit time of day (as temperature conditions were constant) and the assay number as fixed effects.

*Exploration*: The number of hops and flights were fit as the response variable in a Poisson generalized mixed-effect model. On average, we measured individuals 1.23 times in the wild (range: 1 – 4) and 2.89 times in the common garden (range: 1 – 3; approx. 74 to 264 days old; Table S2). For the model on wild birds, we fit age category, Julian date of measure, time of day, and protocol type (old vs. new) as fixed effects. For the CG model, we fit time of day and assay number as fixed effects. We excluded three individuals from this analysis; one individual was injured before the exploration assay (forest female) and two individuals did not regrow all their wing feathers after moulting which affected movement during the exploration assay (one urban female and one forest male).

*Tarsus length:* Tarsus length was fit as the response variable in Gaussian mixed-effect models. On average, we measured individuals 1.7 times in the wild (range: 1 – 8) and 2.92 times in the common garden (range: 1 – 3; approx. 44 to 264 days old; Table S2). As tarsus length is fixed early in life and should not be affected over time, we controlled for fewer confounding effects for this trait. In the wild models, we did not add additional fixed effects. In the CG model, we included observer as a fixed effect, which accounted for both differences between observer measures and differences between life stages as the first observer measured individuals earlier in life and the other observer past 152 days old. We excluded one urban female from this analysis that was a clear outlier for this trait; this individual was very small and undeveloped at 10 days post hatching when entering captivity (e.g., weight = 4.6 g vs. average weight = 13.6 g).

*Body mass*: Body mass was fit as the response variable in Gaussian mixed-effect models. On average, we measured individuals 1.7 times in the wild (range: 1 – 8) and 2.92 times in the common garden (range: 1 – 3; approx. 44 to 264 days old; Table S2). For the wild model, we also controlled for age category, date of measure, and time of day as fixed effects. For the CG model, we also included the time of day and assay number as fixed effects. Besides the individual outlier that we excluded for tarsus length, we also excluded a forest female that was injured before body mass measurements.

*Common garden animal models*

We built upon the common garden models presented in Table 2.2 of the main text by additionally fitting a genetic relatedness matrix (GRM) as a random effect. This approach allowed us to partition variance between the GRM (= individual genetic relatedness; $V_A$), individual ID (= includes individual-specific or permanent environmental variation; $V_{ID}$), and origin nest (= includes differences in maternal investment; $V_{NO}$) random effects, otherwise model structures remained the same. For these animal models, we used weakly informative inverse-Gamma priors (V = 1, nu = 0.002) for fixed effects and parameter-expanded priors (i.e., V = 1, nu = 1, alpha.mu = 0, alpha.V = 1000) for random effects as we expected quantitative genetic parameters to be small (de Villemereuil 2018). We ran all models with MCMCglmm for 1000000 iterations (except tarsus which we ran for 2000000), with a thinning of 500 and a burn-in period of 10000, which achieved effective sample sizes > 1000 across all estimates.

*Heritability and $Q_{ST}$ / $F_{ST}$ comparisons*

We computed the heritability ($h^2$) of each trait in the common garden animal models as:

$$h^2 = \frac{V_A}{V_P} \quad (1)$$

$$\text{where } V_P = V_A + V_{ID} + V_{NO} + V_{NF} + V_{AV} + V_{SEX} + V_{HAB} + V_R \quad (2)$$

where $V_P$ is the total phenotypic variance and comprises variance across individual genetic relatedness ($V_A$), individuals ($V_{ID}$), origin nests ($V_{NO}$), foster nests ($V_{NF}$), aviaries ($V_{AV}$), and residuals ($V_R$). We also included variance generated by non-experimental fixed effects (habitat and sex; $V_{SEX}$ + $V_{HAB}$) in the model (de Villemereuil et al. 2018). For the Poisson animal model (exploration), we used the QCglmm package (de Villemereuil et al. 2016) to convert the variance components and heritability estimate from the latent scale to the data scale.

We also computed $Q_{ST}$ values for the common garden traits where habitat differences were clearly maintained (i.e., breath rate and body mass) so we could make $Q_{ST}$-$F_{ST}$ comparisons and evaluate whether processes other than adaptive evolution, specifically genetic drift, can partly explain phenotypic and genetic differentiation observed between populations (Leinonen et al. 2013). $Q_{ST}$, measured at the phenotypic level in the common garden, quantifies the additive genetic variation between populations relative to the total genetic variance in a phenotype. As we use animal models that directly estimated the additive genetic variation for each trait in the common garden, we estimated $Q_{ST}$ as:

$$Q_{ST} = \frac{V_B}{V_B + 2V_A} \tag{3}$$

where $V_B$ is the between-population genetic variance (habitat fixed effect variance) and $V_A$ is the within-population genetic variance. We additionally corrected $V_B$ by the uncertainty around estimating this effect (product of the beta and design covariance matrices) due to our small sample size. We compared the $Q_{ST}$ value to $F_{ST}$, measured at the molecular level across wild populations, which quantifies neutral molecular variance and represents a null expectation that the observed differentiation between populations is a result of genetic drift and migration. We compared to previous estimates of $F_{ST}$ for the study populations where all urban and forest comparisons were between 0.006 – 0.009 (Perrier et al. 2018). $Q_{ST} > F_{ST}$ (here 0.06 and 0.08 for breath rate and body mass, respectively, Table S4) reveals opposite directional selection pressures in the two populations favouring local adaptation and a stronger divergence in the trait between these populations than expected with genetic drift alone (Leinonen et al. 2013).

**Table S1** Summary of the transfer of eggs to foster nests and nestlings transferred to the nursery under common garden conditions from urban and rural origin habitats in Montpellier, France. Origin nests were spread across four urban sites (FON, MAS, FAC, CEF) and one forest site (ROU), and were transferred to wild foster nests at a single site at the Montpellier Zoo (ZOO). We could not identify the exact origin nest for foster nest ZOO46 as we were missing parental identity and genotyping revealed that these origin nests (FAC2, CEF7) included 3 pairs of siblings.

| Origin habitat | Origin nest ID | Origin lay date (days since Jan 1) | Eggs | Foster nest ID | Average egg weight (g) | Nestlings | Nursery date (days since Jan 1) |
|---|---|---|---|---|---|---|---|
| Urban | FON16 | 91 | 4 | ZOO42 | NA | 4 | 119 |
| | MAS31 | 90 | 4 | ZOO42 | NA | 4 | |
| | FON18 | 91 | 4 | ZOO63 | NA | 4 | 123 |
| | MAS28 | 91 | 4 | ZOO63 | NA | 3 | |
| | FAC7 | 94 | 3 | ZOO69 | 1.49 | 1 | 123 |
| | FAC17 | 94 | 3 | ZOO69 | 1.63 | 3 | |
| | FAC4 | 98 | 3 | ZOO32 | 1.56 | 1 | 126 |
| | CEF9 | 100 | 3 | ZOO32 | 1.33 | 3 | |
| | FON3 | 91 | 4 | ZOO40 | 1.48 | 3 | 126 |
| | MAS40 | 90 | 3 | ZOO40 | 1.57 | 1 | |
| | FAC2 | 101 | 3 | ZOO46 | 1.66 | 2 or 4 | 131 |
| | CEF7 | 100 | 4 | ZOO46 | 1.43 | 2 or 4 | |
| | MAS37 | 107 | 4 | ZOO30 | 1.70 | 4 | 135 |
| | FON5 | 108 | 4 | ZOO30 | 1.74 | 4 | |
| **Total/ Average** | **14** | **97.08** | **50** | **7** | **1.56** | **41** | **126.33** |
| Forest | ROU13s | 98 | 4 | ZOO19 | 1.83 | 4 | 128 |
| | ROU334 | 99 | 4 | ZOO19 | 1.64 | 4 | |
| | ROU12s | 105 | 4 | ZOO67 | 1.66 | 4 | 133 |
| | ROU17s | 104 | 4 | ZOO67 | 1.75 | 4 | |
| | ROU15s | 105 | 4 | ZOO35 | 1.81 | 4 | 131 |
| | ROU9s | 104 | 4 | ZOO35 | 1.67 | 4 | |
| | ROU324 | 108 | 4 | ZOO65 | 1.69 | 0 | NA |
| | ROU7s | 108 | 4 | ZOO65 | 1.62 | 0 | |
| | ROU3s | 109 | 4 | ZOO47 | 1.65 | 4 | 136 |
| | ROU10s | 107 | 4 | ZOO47 | 1.67 | 4 | |
| **Total/ Average** | **10** | **104.70** | **40** | **5** | **1.70** | **32** | **131.86** |

**Table S2** Overview of phenotyping in the common garden experiment including number of observations, individuals, and mean and range of number of repeated measures per individual. The age when individuals were measured is also shown. Note that genotypic data was only available for 72 individuals and so animal models will exclude the one individual that wasn't sequenced.

| Phenotype | Observations | Individuals | Mean number of repeated measures | Range of number of repeated measures | Mean age when assayed (days old) |
|---|---|---|---|---|---|
| Tarsus | 211 | 72 | 2.92 | 1 - 3 | 44.67, 159.25, 264.83 |
| Mass | 210 | 71 | 2.92 | 1 - 3 | 44.67, 159.25, 264.83 |
| Aggression | 282 | 73 | 3.89 | 1 - 4 | 44.67, 74.38, 159.25, 264.83 |
| Breath rate | 284 | 73 | 3.89 | 1 - 4 | 44.67, 74.38, 159.25, 264.83 |
| Exploration | 203 | 70 | 2.89 | 1 - 3 | 74.38, 159.25, 264.83 |

**Table S3** Common garden animal model comparison to Table 3 when including the genetic relatedness matrix used to additionally estimate individual genetic variance ($V_A$). Fixed and random model estimates and 95% credible intervals (CI) across phenotypic traits (A: Aggression in hand, B: Breath rate index, C: Exploration, D: Tarsus length, and E: body mass). Exploration estimates are from a Poisson generalized mixed-effect model, while all other traits were fit with Gaussian mixed-effect models. Interactions between sex and habitat were not significant across traits and dropped from the model. The number of observations (obs) and individuals (ind) for each trait and context are shown in the top panel. We were missing genetic data on one individual and so our sample sizes and observations differ slightly from those reported in Table 3. We also report estimated heritability across all traits and $Q_{ST}$ values for traits where we see a habitat difference (breath rate index and body mass).

| | A) Aggression N = 280 obs, 72 ind | | B) Breath rate N = 279 obs, 72 ind | | C) Exploration N = 200 obs, 69 ind | | D) Tarsus length N = 208 obs, 71 ind | | E) Body mass N = 207 obs, 70 ind | |
|---|---|---|---|---|---|---|---|---|---|---|
| *Fixed effects* | Est | CI | Est | CI | Est | CI | Est | CI | Est | CI |
| Intercept | 1.36 | 0.13 - 2.5 | 9.41 | 7.09 - 11.73 | 4.03 | 1.89 - 6.11 | 19.41 | 19.04 - 19.8 | 15.93 | 14.76 - 17.13 |
| Habitat (urban) | 0.18 | -0.24 - 0.58 | -1.07 | -2.57 - 0.44 | -0.05 | -1.12 - 1.23 | -0.22 | -0.75 - 0.25 | -0.51 | -1.14 - 0.03 |
| Sex (male) | -0.24 | -0.52 - 0.04 | 0.78 | 0.08 - 1.52 | -0.41 | -0.97 - 0.08 | 0.51 | 0.28 - 0.71 | 1.03 | 0.77 - 1.31 |
| Time of day | 0.04 | -0.07 - 0.16 | 0.47 | 0.27 - 0.67 | -0.06 | -0.26 - 0.11 | | | -0.11 | -0.22 - 0 |
| Measurement (2) | -0.17 | -0.47 - 0.1 | 0.88 | 0.43 - 1.31 | -0.19 | -0.62 - 0.27 | | | 0.12 | -0.02 - 0.32 |
| Measurement (3) | -0.25 | -0.52 – 0.003 | 0.45 | -0.01 - 0.84 | -0.09 | -0.53 - 0.35 | | | 0.51 | 0.36 - 0.66 |
| Measurement (4) | -0.40 | -0.63 - -0.13 | -0.40 | -0.77 - 0.03 | | | | | | |
| Observer (2) | | | | | | | 0.07 | 0.04 - 0.1 | | |
| *Random effects* | | | | | | | | | | |
| GRM ($V_A$) | 0.07 | <0.001 - 0.24 | 1.53 | <0.001 - 3.51 | 0.43 | <0.001 - 1.08 | 0.16 | <0.001 - 0.34 | 0.26 | <0.001 - 0.49 |
| Individual ID ($V_{ID}$) | 0.13 | <0.001 - 0.28 | 1.14 | <0.001 - 2.47 | 0.28 | <0.001 - 0.79 | 0.09 | <0.001 - 0.21 | 0.09 | <0.001 - 0.26 |
| Origin nest ID ($V_{NO}$) | 0.02 | <0.001 - 0.09 | 0.48 | <0.001 - 1.62 | 0.22 | <0.001 - 0.74 | 0.08 | <0.001 - 0.24 | 0.07 | <0.001 - 0.25 |
| Foster nest ID ($V_{NF}$) | 0.02 | <0.001 - 0.09 | 0.34 | <0.001 - 1.30 | 0.47 | <0.001 - 1.68 | 0.05 | <0.001 - 0.19 | 0.06 | <0.001 - 0.25 |
| Aviary ID ($V_{AV}$) | 0.02 | <0.001 - 0.10 | 0.48 | <0.001 - 1.75 | 0.13 | <0.001 - 0.51 | | | | |
| Residual variance ($V_R$) | 0.56 | 0.47 - 0.69 | 1.53 | 1.25 - 1.86 | 1.59 | 1.16 - 2 | 0.01 | 0.01 - 0.02 | 0.18 | 0.14 - 0.23 |
| $h^2$ | 0.05 | <0.001 – 0.28 | 0.23 | 0.002 – 0.57 | 0.02 | <0.001 – 0.06 | 0.33 | 0.003 – 0.72 | 0.28 | 0.01 – 0.49 |
| $Q_{ST}$ | | | 0.06 | <0.001 – 0.99 | | | | | 0.08 | <0.001 – 0.82 |

**Table S4** Model outputs for comparison to Table 3 when replacing the habitat effect with the proportion ISA (impervious surface area at 100m; continuous urbanization). Fixed and random model estimates and 95% credible intervals (CI) for 1) wild and 2) common garden contexts across phenotypic traits (A: Aggression in hand, B: Breath rate index, C: Exploration, D: Tarsus length, and E: Body mass). Exploration estimates are from Poisson generalized mixed-effects model, while all other traits were fit with Gaussian mixed-effects models. The number of observations (obs) and individuals (ind) for each trait and context are shown in the top panel.

1) WILD

| | A) Aggression N = 1308 obs, 773 ind | | B) Breath rate N = 702 obs, 531 ind | | C) Exploration N = 581 obs, 472 ind | | D) Tarsus length N = 1474 obs, 861 ind | | E) Body mass N = 1391 obs, 817 ind | |
|---|---|---|---|---|---|---|---|---|---|---|
| *Fixed effects* | Est | CI | Est | CI | Est | CI | Est | CI | Est | CI |
| Intercept | 2.36 | 1.81 - 2.96 | 12.81 | 10.4 - 14.89 | 3.87 | 1.05 - 6.71 | 19.33 | 19.17 - 19.47 | 15.86 | 15.39 - 16.3 |
| ISA | 0.01 | -0.29 - 0.32 | -0.86 | -1.58 - -0.03 | 1.23 | -0.07 - 2.15 | -0.20 | -0.34 - -0.06 | -0.35 | -0.62 - -0.02 |
| Sex (male) | 0.17 | -0.03 - 0.33 | 0.06 | -0.3 - 0.41 | -0.04 | -0.45 - 0.36 | 0.55 | 0.47 - 0.62 | 0.62 | 0.51 - 0.72 |
| Age (yearling) | -0.08 | -0.18 - 0.03 | 0.00 | -0.31 - 0.3 | 0.11 | -0.25 - 0.45 | | | -0.32 | -0.39 - -0.23 |
| Time of day | -0.04 | -0.06 - -0.01 | | | -0.01 | -0.1 - 0.07 | | | 0.04 | 0.02 - 0.06 |
| Date of measure | <0.001 | <0.001 - <0.001 | 0.00 | -0.02 - 0.01 | -0.01 | -0.03 - 0.01 | | | <0.001 | <0.001 - <0.001 |
| Protocol (old) | | | 0.18 | -0.35 - 0.73 | 0.17 | -0.39 - 0.71 | | | | |
| Temperature | | | 0.09 | 0.06 - 0.13 | | | | | | |
| ISA * Sex | 0.26 | <0.001 - 0.57 | | | | | | | | |
| *Random effects* | | | | | | | | | | |
| Individual ID ($V_{IND}$) | 0.45 | 0.36 - 0.54 | 2.63 | 2.07 - 3.18 | 2.83 | 2.08 - 3.67 | 0.28 | 0.25 - 0.31 | 0.38 | 0.33 - 0.45 |
| Site ID ($V_{SITE}$) | 0.02 | <0.001 - 0.08 | 0.24 | <0.001 - 0.76 | 0.21 | <0.001 - 1.08 | 0.01 | <0.001 - 0.04 | 0.03 | <0.001 - 0.13 |
| Year ID ($V_{YEAR}$) | 0.02 | <0.001 - 0.07 | 0.05 | <0.001 - 0.18 | 0.09 | <0.001 - 0.33 | 0.00 | <0.001 - <0.001 | 0.06 | 0.01 - 0.14 |
| Observer ID ($V_{OBS}$) | 0.04 | 0.01 - 0.09 | 0.57 | 0.08 - 1.39 | | | 0.01 | <0.001 - 0.01 | | |
| Residual variance | 0.54 | 0.48 - 0.61 | 1.75 | 1.41 - 2.14 | 1.61 | 1.12 - 2.16 | 0.02 | 0.02 - 0.02 | 0.28 | 0.24 - 0.31 |

2) COMMON GARDEN

| | A) Aggression N = 280 obs, 72 ind | | B) Breath rate N = 279 obs, 72 ind | | C) Exploration N = 200 obs, 69 ind | | D) Tarsus length N = 208 obs, 71 ind | | E) Body mass N = 207 obs, 70 ind | |
|---|---|---|---|---|---|---|---|---|---|---|
| *Fixed effects* | Est | CI | Est | CI | Est | CI | Est | CI | Est | CI |
| Intercept | 1.43 | 0.34 - 2.7 | 9.06 | 6.85 - 11.44 | 3.86 | 1.91 - 6.02 | 19.44 | 19.14 - 19.73 | 15.81 | 14.47 - 16.91 |
| ISA | 0.10 | -0.27 - 0.51 | -0.58 | -2.14 - 1.01 | 0.47 | -0.44 - 1.41 | -0.29 | -0.75 - 0.13 | -0.54 | -1.11 - -0.05 |
| Sex (male) | -0.22 | -0.48 - 0.06 | 0.80 | 0.03 - 1.58 | -0.35 | -0.86 - 0.18 | 0.45 | 0.23 - 0.67 | 1.02 | 0.78 - 1.34 |
| Time of day | 0.04 | -0.08 - 0.15 | 0.46 | 0.26 - 0.67 | -0.06 | -0.25 - 0.14 | | | -0.10 | -0.22 - 0.01 |
| Measurement (2) | -0.17 | -0.46 - 0.11 | 0.94 | 0.48 - 1.37 | -0.19 | -0.62 - 0.31 | | | 0.12 | -0.07 - 0.27 |
| Measurement (3) | -0.26 | -0.5 - <0.001 | 0.49 | 0.08 - 0.95 | -0.13 | -0.57 - 0.34 | | | 0.50 | 0.37 - 0.66 |
| Measurement (4) | -0.39 | -0.64 - -0.14 | -0.38 | -0.79 - 0.04 | | | | | | |
| Observer (2) | | | | | | | 0.07 | 0.04 - 0.11 | | |
| *Random effects* | | | | | | | | | | |
| Individual ID ($V_{ID}$) | 0.19 | 0.08 - 0.32 | 2.11 | 1.11 - 3.25 | 0.49 | <0.001 - 0.97 | 0.18 | 0.11 - 0.27 | 0.27 | 0.14 - 0.42 |
| Origin nest ID ($V_{NO}$) | 0.02 | <0.001 - 0.06 | 0.85 | <0.001 - 2.16 | 0.19 | <0.001 - 0.66 | 0.10 | <0.001 - 0.24 | 0.11 | <0.001 - 0.29 |
| Foster nest ID ($V_{NF}$) | 0.01 | <0.001 - 0.05 | 0.24 | <0.001 - 1.12 | 0.18 | <0.001 - 0.74 | 0.02 | <0.001 - 0.09 | 0.04 | <0.001 - 0.14 |
| Aviary ID ($V_{AV}$) | 0.01 | <0.001 - 0.05 | 0.14 | <0.001 - 0.66 | 0.06 | <0.001 - 0.24 | | | | |
| Residual variance ($V_R$) | 0.56 | 0.45 - 0.67 | 1.58 | 1.3 - 1.91 | 1.65 | 1.18 - 2.1 | 0.01 | 0.01 - 0.02 | 0.18 | 0.14 - 0.22 |

**Table S5** Model outputs for comparison to Table 3.2A showing weak and unclear interaction between habitat and sex on aggression in hand measured in the common garden context; an interaction which was statistically clear for aggression in hand in the wild context. Fixed and random model estimates and 95% credible intervals (CI) are shown and model was fit with a Gaussian mixed-effects model. The number of observations (obs) and individuals (ind) are shown in the top panel.

| | A) Aggression N = 280 obs, 72 ind | |
|---|---|---|
| *Fixed effects* | Est | CI |
| Intercept | 1.31 | -0.10 - 238 |
| Habitat (urban) | 0.24 | -0.21 - 0.66 |
| Sex (male) | -0.08 | -0.48 - 0.32 |
| Time of day | 0.04 | -0.07 - 0.16 |
| Measurement (2) | -0.17 | -0.45 - 0.11 |
| Measurement (3) | -0.25 | -0.52 – 0.02 |
| Measurement (4) | -0.39 | -0.62 - -0.13 |
| Habitat*Sex | -0.25 | -0.76 – 0.32 |
| *Random effects* | | |
| Individual ID ($V_{ID}$) | 0.19 | 0.07 - 0.32 |
| Origin nest ID ($V_{NO}$) | 0.02 | <0.001 - 0.06 |
| Foster nest ID ($V_{NF}$) | 0.02 | <0.001 - 0.06 |
| Aviary ID ($V_{AV}$) | 0.01 | <0.001 - 0.05 |
| Residual variance ($V_R$) | 0.56 | 0.46 - 0.67 |

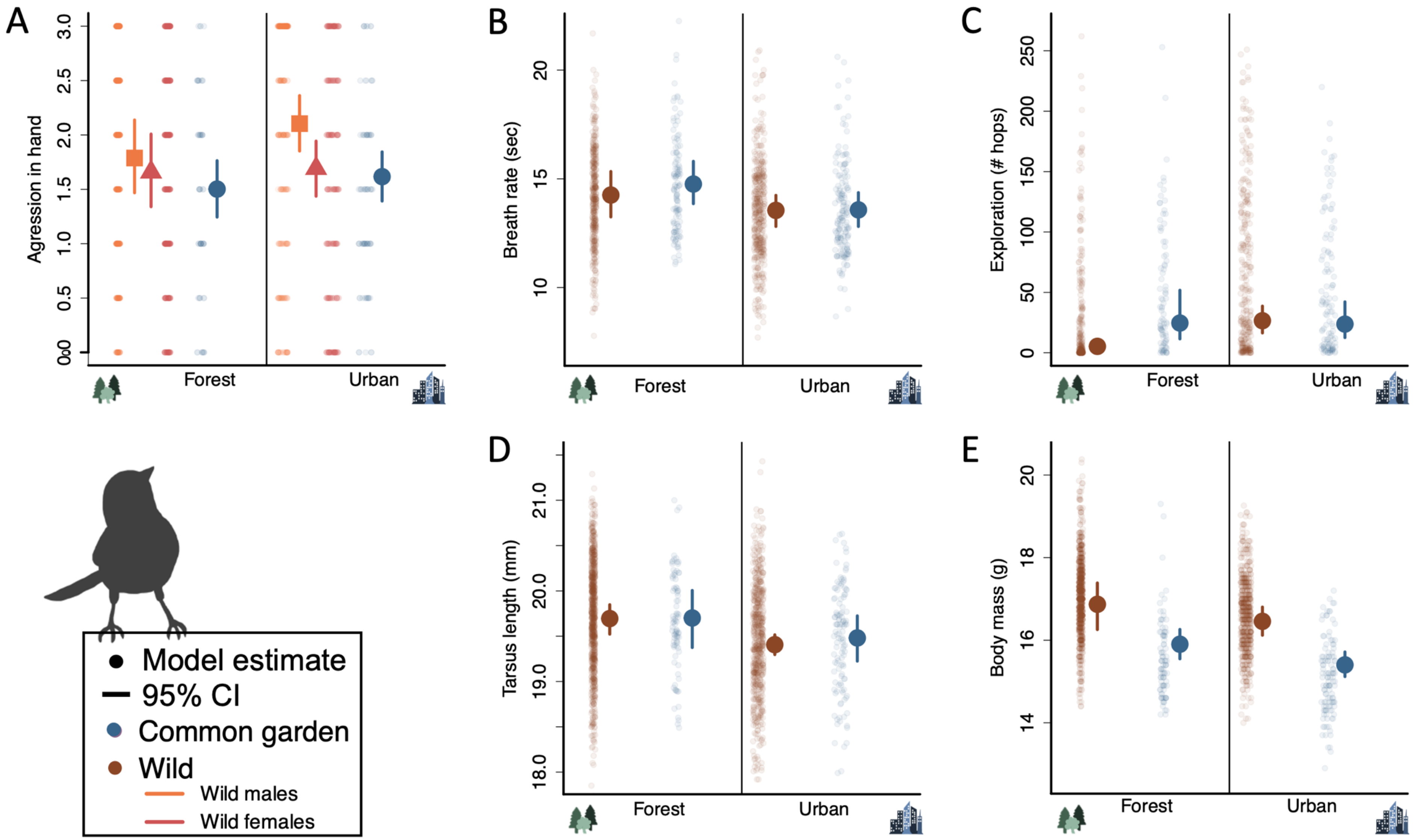


**Figure S1.** Recreation of Figure 1 in text showing model effects with full distribution of raw data. Habitat type model estimate and 95% credible intervals (CI) on phenotypic traits of wild (brown) and common garden birds (blue) across A) aggression in hand, B) breath rate index, C) exploration score, D) tarsus length, and E) body mass. Habitat differences varied clearly by sex only in one case (A: aggression in wild birds) and these sex differences are shown (wild males: squares, wild females: triangles); aggression did not differ clearly by sex in the common garden. Note that random noise around raw data has been added to better visualize overlapping points.

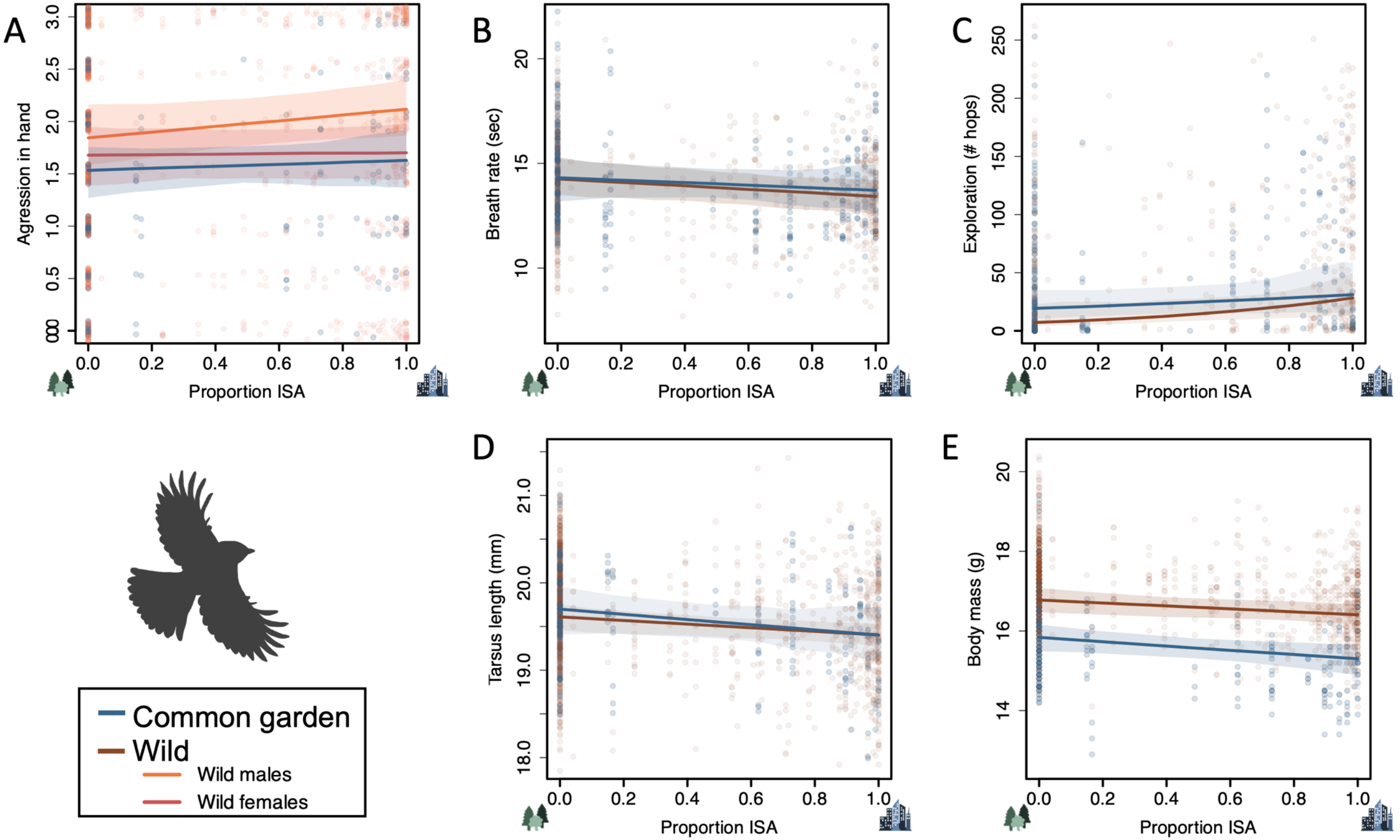


**Figure S2.** Recreation of Figure 1 in text showing model effects with full distribution of raw data. Proportion of impervious surface area **(**ISA) model estimate and 95% credible intervals (CI) on phenotypic traits of wild (brown) and common garden (blue) birds across A) aggression in hand, B) breath rate index, C) exploration score, D) tarsus length, and E) body mass. ISA effects varied clearly by sex only in one case (A: aggression in wild birds) and these sex differences are shown (wild males: orange, wild females: red); aggression over the ISA gradient in the common garden did not differ clearly by sex. Note that random noise around raw data has been added to better visualize overlapping points.

**References:**


Caizergues AE, Grégoire A, Choquet R, Perret S, Charmantier A. 2022. Are behaviour and stress-related phenotypes in urban birds adaptive? Journal of Animal Ecology. 91(8):1627–1641.

European Environment Agency. 2020. Imperviousness Density 2018. https://land.copernicus.eu/user-corner/technical-library/imperviousness-2018-user-manual.pdf.

Huisman J. 2017. Pedigree reconstruction from SNP data: parentage assignment, sibship clustering and beyond. Molecular Ecology Resources. 17(5):1009–1024. doi:10.1111/1755-0998.12665.

Leinonen T, Mccairns RJS, Hara RBO, Merilä J. 2013. QST – FST comparisons : evolutionary and ecological insights from genomic heterogeneity. Nature Reviews Genetics. doi:10.1038/nrg3395.

Perrier C, Lozano del Campo A, Szulkin M, Demeyrier V, Gregoire A, Charmantier A. 2018. Great tits and the city: Distribution of genomic diversity and gene–environment associations along an urbanization gradient. Evolutionary Applications. 11(5):593–613. doi:10.1111/eva.12580.

QGIS Development Team. 2023. QGIS Geographic Information System. https://www.qgis.org.

de Villemereuil P. 2018. Quantitative genetic methods depending on the nature of the phenotypic trait. Annals of the New York Academy of Sciences. 1422(1):29–47. doi:10.1111/nyas.13571.

de Villemereuil P, Morrissey MB, Nakagawa S, Schielzeth H. 2018. Fixed-effect variance and the estimation of repeatabilities and heritabilities: Issues and solutions. Journal of Evolutionary Biology. 31(4):621–632.

de Villemereuil P, Schielzeth H, Nakagawa S, Morrissey M. 2016. General methods for evolutionary quantitative genetic inference from generalized mixed models. Genetics. 204(3):1281–1294. doi:10.1534/genetics.115.186536.
